# Thirty years of *Achromobacter ruhlandii* evolution reveal pathways to epidemic lineages

**DOI:** 10.64898/2026.03.25.714254

**Authors:** Migle Gabrielaite, Helle Krogh Johansen, Jonas Juozapaitis, Rasmus L. Marvig, Gytis Dudas

## Abstract

**Background:** *Achromobacter* spp. are emerging opportunistic pathogens, associated with chronic infections, antimicrobial resistance, and poor clinical outcomes. The Danish epidemic strain (DES) of *A. ruhlandii* is highly drug-resistant and adapted to the cystic fibrosis (CF) airway, yet its evolutionary history and defining genomic features remain poorly understood.

**Methods:** We analysed genome and antibiotic susceptibility testing data for 58 longitudinally collected DES isolates sampled over 21 years at Rigshospitalet, Denmark. We combined these with 79 publicly available *A. ruhlandii* genomes and applied phylogenomics to infer DES emergence and transmission, and genome-wide association studies (GWAS) to identify lineage-specific and adaptive genomic features.

**Results:** DES forms a distinct monophyletic clade within *A. ruhlandii*, estimated to have emerged around 1990, with no evidence of dissemination beyond Denmark. GWAS identified key lineage-defining features, including acquisition of large mobile genetic elements and enrichment of resistance and iron acquisition genes. In addition, we detected other epidemic *A. ruhlandii* lineages with evidence of long-term persistence and inter-country spread, sharing similar genetic signatures of adaptation.

**Conclusions:** This study elucidates the genomic features associated with chronic infection and epidemic potential in *A. ruhlandii*. The DES lineage exemplifies how extensive horizontal gene transfer, high intrinsic resistance potential, and host-adaptation traits, such as increased iron acquisition, can facilitate the emergence and persistence of successful epidemic lineages. These findings highlight shared evolutionary signatures of epidemic *A. ruhlandii* and underscore the need for continued genomic surveillance to detect and monitor emerging high-risk lineages in chronic infections.

## Introduction

*Achromobacter* spp. are emerging opportunistic pathogens with natural reservoirs in soil and water ^1^. In clinical settings, they are increasingly associated with chronic airway infections in immunocompromised individuals, particularly people with cystic fibrosis (pwCF) with comparable risk of disease deterioration as in *Pseudomonas aeruginosa* infections ^2^.

*Achromobacter* is reported to account for up to 20% of respiratory infections in pwCF in Europe in 2023 ^3^, markedly higher than the 2018 estimate of less than 10% ^4^. Notably, *Achromobacter* spp. prevalence is likely underestimated in many countries due to identification difficulties and common misclassification as *P. aeruginosa*, *Stenotrophomonas maltophilia* or *Burkholderia cepacia* complex ^1,5,6^.

To date, twenty-two different *Achromobacter* species have been described in NCBI and LSPN ^7,8^. However, taxonomic boundaries within the genus remain unclear, with more than 10% of isolates misidentified even in whole-genome sequencing (WGS)-based analyses ^9^. The Genome Taxonomy Database (GTDB) currently recognises 55 *Achromobacter* species ^10^, underscoring extensive genomic diversity within the genus and the recent delineation of several new species that were previously grouped under more common taxa such as *A. xylosoxidans, A. insuavis*, and *A. spanius*. Despite this diversity, research has predominantly focused on *A. xylosoxidans*, which accounts for over half of all clinical infections, while less abundant species such as *A. ruhlandii* have received comparatively limited attention ^2,5,11^.

Clinical *Achromobacter* spp. isolates are highly resistant to antibiotics, reflecting both intrinsic and acquired resistance mechanisms ^12^. These include multidrug efflux systems such as AxyXY-OprZ and AxyABM, β-lactamases, and other genes (e.g., *aac(6*′*)Ib-cr*, *qnrA*, *oqxA*, *oqxB*), often acquired through horizontal gene transfer, conferring commonly observed resistance to β-lactams, fluoroquinolones, aminoglycosides, and even carbapenems ^1,13,14^. This multidrug-resistant (MDR) phenotype complicates eradication and contributes to the persistence of *Achromobacter* spp. in the airways despite intensive antimicrobial treatment ^1^.

Cystic Fibrosis Transmembrane Regulator protein (CFTR) modulator therapy has reduced the overall prevalence of airway infections in pwCF ^15,16^; nonetheless, multiple studies have reported shifts in the lung microbiome, suggesting alterations in the dynamics of bacterial colonisation and infection ^17,18^. Of particular concern is the Danish Epidemic Strain (DES): a lineage of *Achromobacter ruhlandii* that is highly prevalent among pwCF in Denmark, exhibits high level of resistance to antibiotics, a hypermutator phenotype, and continues to be transmitted between patients since at least 1993 ^14,19,20^.

Despite its clinical importance, the evolutionary history, origin, and genomic features of DES remain poorly understood. Here, we use 158 whole-genome sequenced isolates of *A. ruhlandii* to perform comparative genomics, phylogenetic analysis, and genome-wide association study (GWAS) to investigate the emergence, diversification, and adaptation of DES. By integrating genomes of longitudinal isolates of DES with publicly available genomes of *A. ruhlandii*, we aim to elucidate the genomic features that enable epidemic success, with implications for infection control and management of chronic bacterial infections. Furthermore, by analysing additional epidemic *A. ruhlandii* lineages, we generalise these findings to reveal shared evolutionary signatures of other epidemic lineages in this species.

## Methods

### Sample collection

Sixty-five *A. ruhlandii* clinical isolates obtained from 19 pwCF attending the Copenhagen Cystic Fibrosis Center at Rigshospitalet, Denmark were analysed. This cohort represents 15% (19 of 124) of all patients with *Achromobacter* spp. infections who had isolates’ genomes sequenced at least once between 2002 and 2024 (Supplementary Figure 1) and 15% (19 of 126) of all pwCF with *Achromobacter* spp. infections during this period. Of the 65 *A. ruhlandii* clinical isolates from Rigshospitalet, 36 were newly sequenced isolates, and 29 genomes were previously published under BioProject PRJEB39108 ^9,14^. The use of clinical isolates was approved by the local ethics committee at the Capital Region of Denmark (Region Hovedstaden; approval registration number H-21078844), and the use of clinical registry data was approved by the Danish Agency for Patient Safety (approval registration number 31-1521-428).

All patients received early antibiotic therapy targeting *Achromobacter* spp. upon the first positive culture, guided by antimicrobial susceptibility testing (AST). The most common regimen consisted of inhaled colistin (CST) combined with amoxicillin–clavulanic acid (AMC) for three weeks. If eradication failed, patients typically received a 14-day course of intravenous piperacillin–tazobactam (TZP), meropenem (MEM), or ceftazidime (CAZ) in combination with tobramycin (TOB) and trimethoprim–sulfamethoxazole (SXT). In selected cases, inhaled or oral CST or CAZ was also administered ^14^.

### Susceptibility testing

Antibiotic susceptibility was detected using Neo-Sensitabs from Rosco Diagnostica antibiotic-containing tablets and corresponding zone-of-inhibition interpretive breakpoints. Manufacturer’s guidelines for susceptibility interpretation was used as no EUCAST criteria for *A. ruhlandii* is available. Isolate susceptibility was classified as resistant (R); susceptible, increased exposure (I); or susceptible (S) according to the manufacturer’s guidelines for *Achromobacter* spp. Susceptibility data were available for all 65 *A. ruhlandii* isolates sequenced in this study (Supplementary Table 1). To further assess longitudinal resistance dynamics in the 14 pwCF infected with DES, AST from 1,542 non-sequenced *Achromobacter* spp. isolates (15–230 per patient) were additionally analysed. Chronic infection was defined as ≥4 positive cultures per year ^21^). The included patients were P0904, P1705, P1903 (from 16 December 2011), P3403 (until 1 January 2007), P3704, P4203 (from 9 December 2014), P5303, P5604, P7003, P7703, P8003, P8703, P9503, and P9603. For each patient, the isolate inclusion timeline was adjusted to exclude periods in which non–*A. ruhlandii* isolates were detected. The analysis included 12 antibiotics for which AST was done to the majority of the isolates: AMC, CAZ, chloramphenicol (CHL), ciprofloxacin (CIP), CST, imipenem (IPM), MEM, TZP, sulfamethizole (SMZ), tigecycline (TGC), trimethoprim (TMP), and SXT. Susceptible, increased exposure (I) isolates were classified as susceptible. For each antibiotic, longitudinal resistance was modelled using a binomial generalised additive mixed model, with resistance status as the binary outcome, sampling date as a penalised smooth term and patient as a random intercept. Population-level resistance probabilities were predicted across the observed sampling period for each antibiotic.

### Short read Illumina sequencing

Genomes were sequenced using either the “Nextera” (17 isolates) or the “Hackflex” (19 isolates) workflows employed in the routine diagnostic work at the Department of Genomic Medicine at Rigshospitalet. In the Nextera workflow, genomic DNA was extracted and purified using a DNeasy Blood and Tissue kit (Cat. 69504, Qiagen). Genomic DNA libraries that were prepared using a Nextera XT DNA Library Prep kit (Cat. Illumina) and sequenced on either a MiSeq or NextSeq 500 instrument, generating 250 or 150 base paired-end sequencing reads, respectively. In the Hackflex workflow, an in-house developed DNA extraction procedure was used. DNA was extracted from overnight growth agar plates and subjected to various incubation steps with lysostaphin, lysozyme, proteinase K, and buffer AL (Cat. 19075, Qiagen). After ethanol precipitation and column-based purification, samples were washed, and DNA was eluted in Nuclease free water by centrifugation. The Hackflex workflow libraries were sequenced on either a NextSeq 500 or NovaSeq 6000 instruments, generating 150 base paired-end sequencing reads. Genomes were sequenced with an average of 2,450,753 read pairs (range, 385,790 to 6,935,710 read pairs).

### Public data

Additional 93 genomes from 13 countries from NCBI SRA and Genbank public databases were included in the analysis ^22^. Of these, 44 were isolated from sputum, 28 from other clinical samples (i.e., isolates obtained from clinical specimens other than sputum, e.g., blood, bronchoalveolar lavage, other respiratory specimens), 16 from environmental sources, and five from unknown origins. Of the 93 additional genomes, 28 assemblies were downloaded from AllTheBacteria (retrieved 2024 October) ^23^; 31 assembled genomes were obtained from NCBI Genbank database (retrieved 2024 October); and 34 samples were downloaded as raw sequencing reads and locally assembled (Supplementary Table 2).

### Long read Nanopore re-sequencing and preprocessing

One genome of an early *A. ruhlandii* isolates from Rigshospitalet (Biosample IDs: SAMEA7025168 sampled in 2004) was resequenced using the Oxford Nanopore platform with R10.4.1 Nanopore flow cell, Native barcoding kit 96 (SQK-NBD114-96), and dorado v0.9.5 for basecalling using sup v5.0.0 model ^24^. Hybrid genome assembly was generated using autocycler v0.5.0 ^25^, that integrates the results of multiple assemblers: Canu v2.3 ^26^, Flye v2.9.6 ^27^, miniasm v0.3-r179 ^28^, NECAT v0.0.1 ^29^, NextDenovo v2.5.2 ^30^, and Raven v1.8.3 ^31^. Assembly was polished with Medaka v2.1.0 ^32^ for error correction, followed by polishing with Illumina short reads using Polypolish v0.6.0 ^33^.

The assembled genome was annotated using Bakta v1.10.0 ^34^ with the full reference database v5.1 and default settings. Functional annotation of predicted coding sequences was performed with eggNOG-mapper v2.1 ^35^ using DIAMOND ^36^ search mode. Annotations were restricted to one-to-one orthologs and with Gene Ontology (GO) terms from all evidence categories included. The resequenced SAMEA7025168 that was completely assembled to a single chromosome was used as a reference genome for all further analyses.

### Search for related genomes in public sequence repositories

Pebblescout ^37^ was employed to identify high-similarity genomes using Metagenomic Volume 1, Metagenomic Volume 2, WGS, RefSeq, and SRA Microbe databases. Searches were performed using default parameters, and genomes exhibiting ≥80% coverage of 25-mers were considered high-similarity matches.

### Whole genome sequencing data preprocessing

Raw sequencing reads from Rigshospitalet and the NCBI databases were assembled using the nf-core/BacAss pipeline v2.3.1 ^38^. Briefly, the pipeline uses fastp ^39^ for quality trimming and Unicycler ^40^ for assembly that employs SPAdes ^41^ for short-read assemblies and integrates miniasm ^28^ and Racon ^42^ for long-read assembly. For long-read assemblies, additional polishing is performed using medaka ^32^. Assembly quality was assessed using BUSCO v5.7.1 ^43^ and QUAST v5.3.0 ^44^, and genomes were excluded if they failed to meet at least one of the following criteria: 1) a BUSCO completeness score above 90% for *Achromobacter* spp., 2) less than 500 contigs longer than 1,000 bp, or 3) a total assembly size within the expected range of 5–7.5 Mbp. In total, 19 genomes were excluded from downstream analyses (five newly sequenced genomes from Rigshospitalet and 14 genomes from the NCBI database) resulting in a final dataset of 139 *Achromobacter* spp. genomes (Supplementary Table 2). All assembled genomes were confirmed to belong to *A. ruhlandii* species with GTDB-Tk v2.4.0 ^45^ using v220 database version and then annotated with bakta v1.10.0 ^34^ using the full database v5.1 and default settings.

### Alignment to the reference genome and variant calling

All isolates were aligned to the long-read–sequenced SAMEA7025168 reference genome, and variants were called using Snippy v4.6.0 ^46^ with thresholds of ≥10× coverage, minimum variant quality of 100, and variant allele frequency of ≥0.7. The resulting consensus alignments were used to construct phylogenetic trees. Gene-level coverage was calculated with mosdepth v0.3.10 ^47^ to generate a gene presence–absence matrix. A gene was considered present when its mean coverage reached at least 25% of the average genome coverage.

### Pan- and core-genome construction

The pangenome for 139 *A. ruhlandii* isolates was constructed with Panaroo v1.5.1 ^48^ using moderate clean mode applying a sequence identity threshold of 0.95 and a maximum length-difference threshold of 0.9. Genes were defined as core if present in ≥95% of isolates, with a maximum allowed entropy of 0.1. Core genome alignments, as defined by Panaroo, were generated using MAFFT v7.525 ^49^.

### Genomic feature of interest identification

Genomic feature identification was performed using multiple tools. Abricate v1.0.1 ^50^ was run with the PlasmidFinder ^51^ database for plasmid identification. A suspected plasmid-like fragment was further examined using PIPdb ^52^. Genomic islands were identified with IslandViewer 4 ^53^ that integrate IslandPick ^54^, IslandPath-DIMOB ^55^ and SIGI-HMM ^56^ tools. Proviral sequences were detected using geNomad v1.8.1 ^57^ with default parameters and database v1.7. Virulence genes were identified by aligning against the complete VFDB database ^58^ using blastn ^22^ with an e-value threshold of 0.001; only hits with ≥70% identity and ≥200 bp length were retained. Iron-acquisition genes were defined based on bakta ^34^ annotations corresponding to *siderophore, ferric, TonB, fecR, fecI, FecCD, FatB, afuA, iron transport, Iron ABC transporter, Fe transport,* or *Fe³ transport functions*, and all candidates were manually verified to belong to iron-acquisition pathways. Resistance genes were characterised using AMRFinderPlus v4.0.23 ^59^ using 50% identity and 50% coverage thresholds. Additionally, putative resistance genes were defined using COG annotation from eggNOG-mapper v2.1 ^35^ that were further explored using CARD database v4.0.1 ^60^ and blastp^22^ with e-value threshold of 10^-30^, 60% query sequence coverage and a minimum of 40% sequence identity. All the putative resistance genes were manually evaluated.

### Phylogenetic analysis

Phylogenetic trees for 1) *A. ruhlandii* core genome, 2) DES core genome and 3) DES reference-based alignment were built using IQ-TREE2 ^61^ under a HKY model with gamma-distributed rate heterogeneity across sites (HKY+G4, four discrete categories) ^62,63^. We assessed recombination using ClonalFrameML v1.13 with the default parameters ^64^. For the joint *A. ruhlandii* phylogeny, trees were rooted using an *A. dolens* outgroup. Following rooting, the *A. dolens* sequence was removed from the displayed tree and excluded from subsequent analyses, while retaining the root position established by the outgroup. The DES phylogeny was rooted using the earliest available DES isolate (SAMEA7025159). For the DES reference-based alignment estimated recombination-to-mutation ratio (r/m) was <0.1, therefore, recombination correction was not performed prior to downstream analyses.

### Most recent common ancestor inference

To estimate the time to the most recent common ancestor (tMRCA) of the Danish epidemic strain (DES), we applied complementary regression- and Bayesian-based phylogenetic dating approaches. Specifically, we used two root-to-tip regression methods (TempEst and TreeTime) to assess temporal signal and obtain initial rate estimates, and two Markov chain Monte Carlo (MCMC)-based frameworks (BEAST and BactDating) to infer evolutionary rates and divergence times while accounting for phylogenetic uncertainty.

Root-to-tip genetic distances were regressed with TempEst v1.5.3 ^65^ against sampling dates using the maximum-likelihood phylogeny inferred from the recombination-masked alignment. The tree root was optimised to maximise the correlation between genetic divergence and sampling time. Divergence times were additionally estimated with TreeTime v0.11.4 ^66^, which applies root-to-tip regression on the same phylogeny to provide an independent, non-parametric estimate of the evolutionary rate and time to the most recent common ancestor (tMRCA).

Bayesian time-scaled phylogenetic analyses were performed in BEAST v10.5.0 ^67,68^ on the DES reference genome-based alignment using a HKY substitution model with gamma-distributed rate heterogeneity across sites, four discrete categories ^62,69^. Sampling dates were used to calibrate the molecular clock. We employed a Hamiltonian Monte Carlo relaxed clock and tested an exponential population size prior ^68,70^. Two MCMC chain runs of 200,000,000 iterations were performed, sampling every 20,000 steps. The first 20,000,000 iterations were discarded as burn-in. Convergence and effective sample sizes (ESS >200) for all parameters were assessed using Tracer v1.7.2 ^71^. Posterior estimates of the evolutionary rate and tMRCA were summarised using 95% highest posterior density (95% HPD) intervals. In addition, tMRCA was independently estimated using BactDating v1.1.3 ^72^, which performs Bayesian dating on a recombination-aware phylogeny. The maximum-likelihood tree inferred from the recombination-aware alignment was used as input, with sampling times provided as tip dates and an additive relaxed molecular clock model applied. Three MCMC chains of 1,000,000 iterations were run to assess convergence, and posterior distributions of the evolutionary rate and node ages were evaluated using 95% HPD intervals. Concordance among estimates obtained using root-to-tip regression, TreeTime, BEAST, and BactDating was used to assess the robustness of the inferred tMRCA. All log and XML files from BEAST runs are available on GitHub: https://github.com/MigleSur/DES_supplementary_files.

### Inference of effective population size and mutation accumulation dynamics

Bayesian inference of changes in effective population size was performed in BEAST v10.5.0 ^67,68^ on the DES reference genome-based alignment using a Skygrid coalescent tree prior with effective population size estimated at 6 evenly spaced time points and a cut-off of 36 years ^73,74^. Analyses were conducted under an HKY substitution model with gamma-distributed rate heterogeneity across sites ^62,69^. MCMC chains were run for 50,000,000 iterations, sampling every 5,000 steps, with the first 5,000,000 iterations discarded as burn-in. Convergence and ESS (ESS >200) for all parameters were assessed using Tracer v1.7.2 ^71^.

In parallel, Bayesian time-scaled phylogenetic analyses assuming a constant effective population size were performed in BEAST v10.5.0 ^67,68^ under the same HKY model with gamma-distributed rate heterogeneity across sites ^62,69^. MCMC settings matched those used for the Skygrid analysis. Convergence and ESS (ESS >200) for all parameters were assessed using Tracer v1.7.2 ^71^.

The dN/dS ratio was approximated by adapting the approach described in Dudas *et al.*, 2021^75^. For in-frame protein-coding sequence alignments, four sample groups (DES, US-ES, ST36 and non-ES) were considered separately. A consensus sequence was generated for each group, and each synonymous and non-synonymous substitution was counted only once to be conservative. The observed numbers of synonymous and non-synonymous substitutions were then normalised by the respective numbers of possible synonymous and non-synonymous substitutions, and the ratio of the resulting dN/dS ratio was calculated for each group. The transition/transversion (Ts/Tv) ratio was obtained from BEAST v10.5.0 ^67,68^ using the same HKY model with gamma-distributed rate heterogeneity across sites ^62,69^, a constant effective population size and a strict molecular clock. MCMC settings matched those used for the Skygrid analysis. Convergence and ESS for all parameters were assessed using Tracer v1.7.2^71^. All log and XML files from BEAST runs are available on GitHub: https://github.com/MigleSur/DES_supplementary_files.

### Genomic feature association with epidemic features in *A. ruhlandii*

Associations between iron acquisition genes, virulence factors, and resistance genes and epidemic phenotypes of the three identified epidemic strains (ES): DES, US-ES, and ST36, were evaluated using Bayesian generalised linear multivariate models implemented in the brms R package v2.23.0 ^76,77^. To account for non-independence due to shared ancestry and repeated sampling, recombination-corrected phylogeny and patient identity were included as independent random effects. For each model, MCMC sampling was performed for 50,000 iterations, with the first 25,000 iterations discarded as burn-in. Seven independent chains were run per model. Convergence was assessed by inspection of trace plots and ESS. Group differences were assessed using Bayesian multivariate models, with uncertainty summarised as 95% highest density regions (HDRs), while accounting for phylogenetic relatedness and repeated sampling from the same patient.

### Genome-wide association analysis of epidemic features in *A. ruhlandii*

We performed genome-wide association analyses using Pyseer v1.4.0 ^78^ to identify genomic features associated with DES and other epidemic strains.

For DES comparison with non-DES isolates, we used the gene presence–absence matrix derived from alignment to the SAMEA7025168 reference genome. To account for population structure, we generated a similarity matrix from the recombination-corrected maximum-likelihood phylogeny and applied a linear mixed model. Bonferroni correction p-value threshold of 4.07×10^-5^ was used to account for multiple testing. Identified associations were queried with MetaGraph ^79^ against 2.7 million AllTheBacteria ^23^ assemblies to identify the organisms of highly similar sequences.

For comparisons across all three epidemic strains, GWAS was performed on the gene presence–absence matrices of the pangenome to identify features distinguishing each epidemic strain (DES, US-ES, and ST36) from non-epidemic isolates. Recombination-masked maximum-likelihood phylogenies were used to control for population structure, and significance was assessed with Bonferroni correction using threshold values of 1.19×10^-5^, 8.12×10^-6^, and 7.87×10^-6^ for DES, US-ES, and ST36, respectively.

## Results

To investigate genomic features associated with the epidemic phenotype in *A. ruhlandii*, we focused on the DES, a lineage previously linked to persistent infections, patient-to-patient transmission, and elevated antibiotic resistance in pwCF ^14,19,20^. We queried our laboratory isolate database and public NCBI database to assemble a genomic dataset comprising DES and non-DES *A. ruhlandii* genomes.

A total of 158 *A. ruhlandii* genomes were assembled for this study, including 93 publicly available genomes from NCBI, including 29 public genomes from Rigshospitalet, and 36 newly sequenced isolates from Rigshospitalet. Nineteen genomes did not meet the predefined quality control (QC) thresholds (five from Rigshospitalet isolates and fourteen from public datasets) and were excluded from downstream analyses. The remaining 139 genomes, comprising 60 DES and 79 non-DES *A. ruhlandii* isolates, were retained (Figure 1; Supplementary Figure 2). In addition to DES, two additional ES were identified: US-ES circulating in multiple states of the U.S. sampled over nine years and ST36 ES circulating in the U.S., Russia, and United Kingdom sampled over 14 years (Supplementary Table 2). In this study, an epidemic strain is defined as a genetically distinct bacterial lineage that undergoes sustained transmission across individuals or populations, evidenced by multi-year, multi-region detection and monophyletic clustering in the core-genome phylogeny. We combined genomic, phylogenetic, and GWAS analyses to investigate the emergence and evolutionary trajectory of DES, characterise its genetic distinctiveness, and identify genomic features associated with its epidemic phenotype.

**Figure 1.**
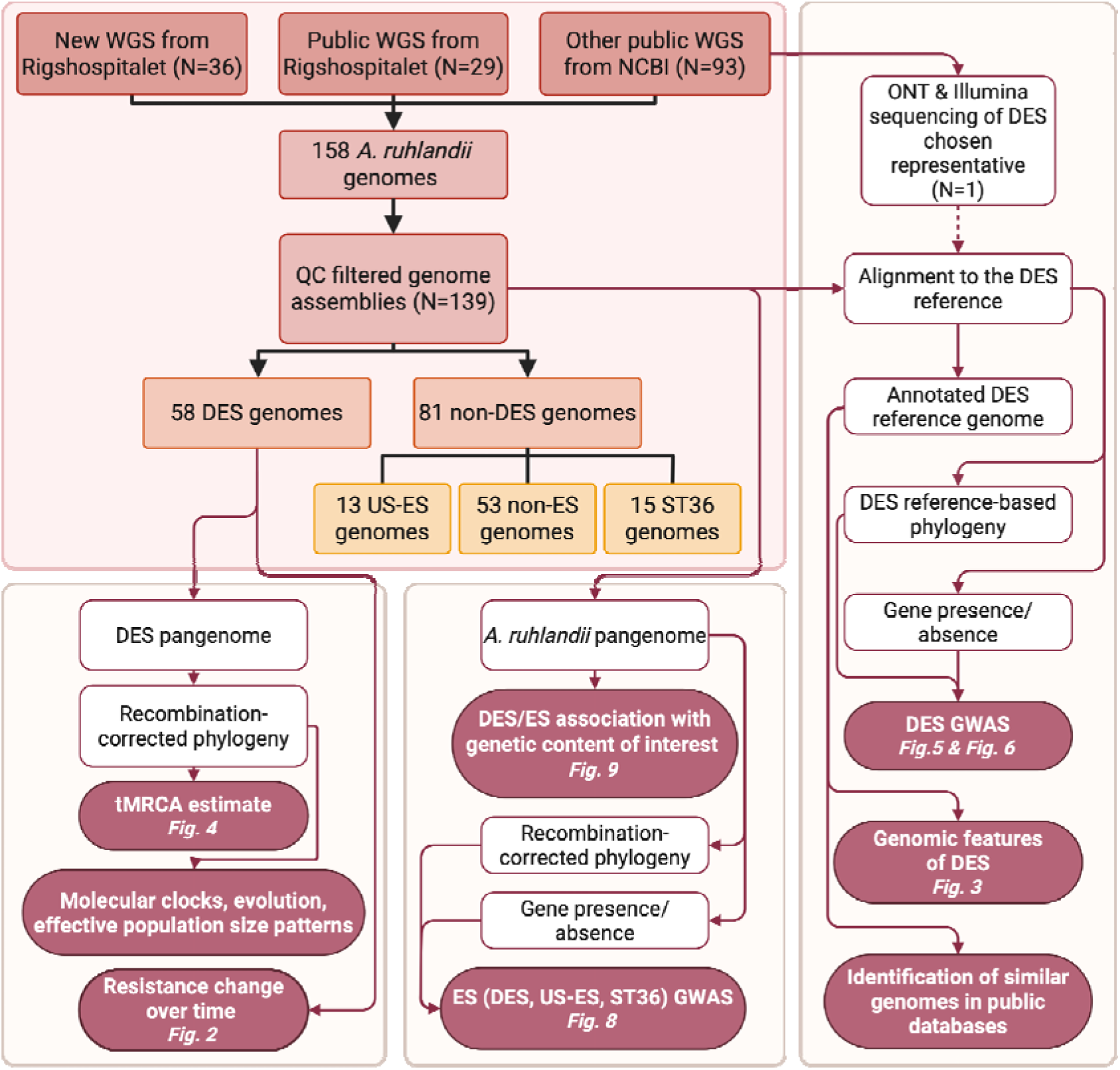
Schematic visualisation of the data set and all the analyses performed in this study.

### Danish epidemic strain is restricted to Denmark

During MLST screening using the DES sequence type ST385 from the PubMLST database ^80^ we identified a single isolate outside Denmark, initially reported from Belgium (ID: 786) that was later confirmed to have been sampled in Denmark ^19^. A metagenomic similarity search using Pebblescout ^37^, indexing both metagenomic and WGS datasets, revealed no high-identity matches (≥80% coverage of 25-mers) to DES. Together, these findings indicate that DES has not been detected outside Denmark.

### DES susceptibility to antibiotics is increasing

We assessed antimicrobial susceptibility of DES isolates collected during clinical visits (N=15–230 isolates per pwCF infected with DES). DES showed high baseline resistance in early isolates for TMP, CAZ, CHL, CIP, SMZ, and TGC (Supplementary Figure 3). Patient-level resistance patterns varied over time, whereas population-level generalised additive mixed models identified increasing resistance over time with decrease in resistance in the recent years for CAZ, CST, IPM, MEM, and SMZ (Figure 2; Supplementary Table 3).

**Figure 2.**
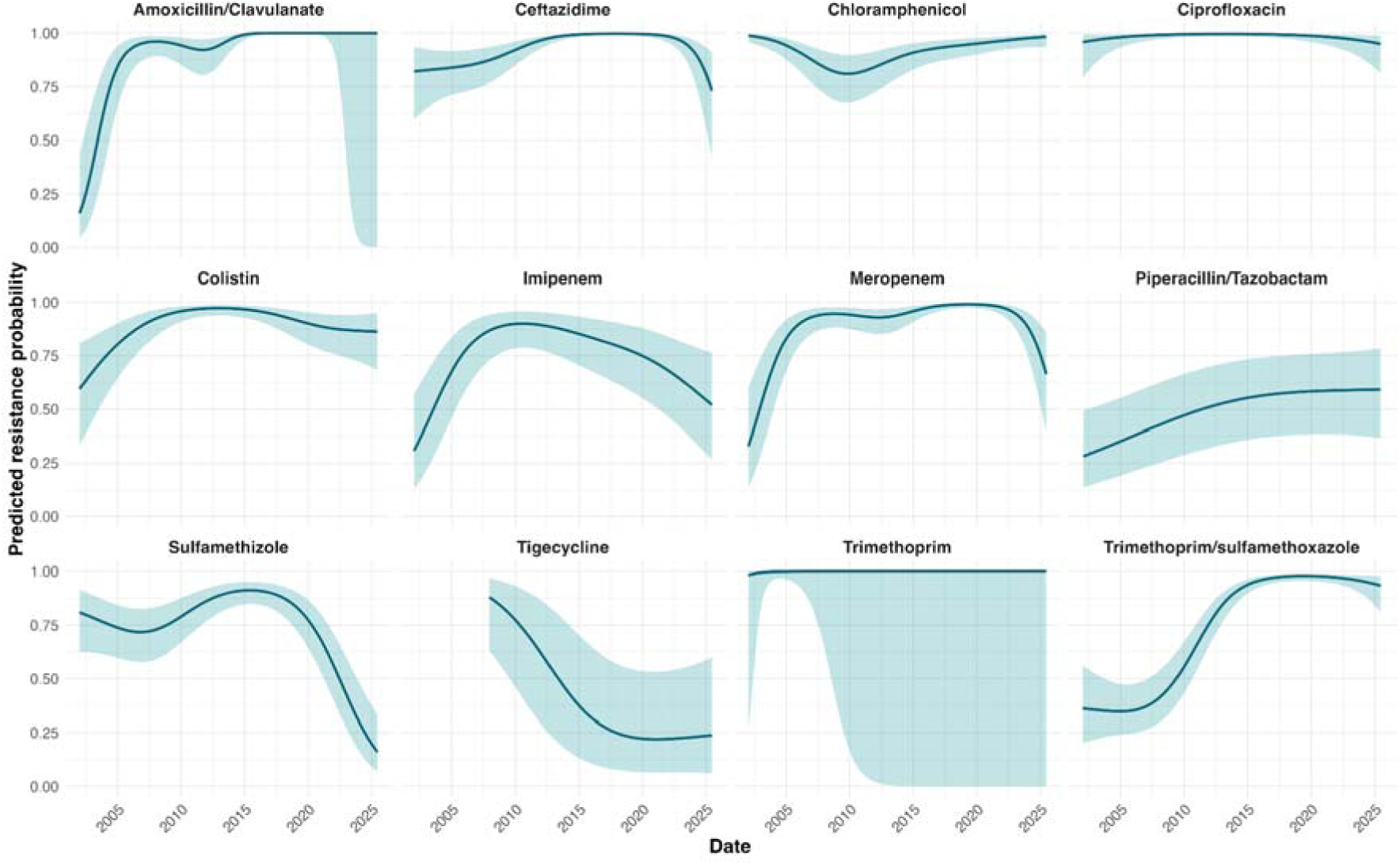
Antibiotic-specific temporal trends in resistance. Predicted probabilities of resistance over time for each antibiotic were estimated using binomial generalised additive mixed-effects models with patient-specific random effects. Solid lines show population-level fitted trends and shaded bands indicate 95% confidence intervals.

### DES resistance cannot be fully explained by current knowledge

AMRFinderPlus identified six recognised resistance genes in the reference DES genome (*bla_AXC_*, *bla_OXA_*, *bla_ELC_*, *tet(A)*, *catB10*, *tmexD*) conferring resistance to β-lactams (including carbapenems) tetracyclines, and phenicols (Supplementary Table 4). All six genes were present in all DES isolates; however, the observed resistance of 95–100% of isolates to Ampicillin, Aztreonam, Ceftriaxone, Cefuroxime, Moxifloxacin, Penicillin, Tetracycline, Tobramycin, TMP, Ceftazidime/Avibactam, and Ceftolozane/Tazobactam antibiotics cannot be explained by these genes alone (Supplementary Table 1). Beyond these known resistance determinants, we identified 59 defense-related genes (COG V) using the DES reference genome, of which 17 were supported as high-confidence putative resistance genes based on CARD orthology and overlapped with three out of six genes identified by AMRFinderPlus (*bla_AXC_*, *bla_OXA_*, *bla_ELC_*). Through manual annotation of the reference DES genome, we defined the additional list of 45 putative resistance-associated genes. Of these, 41 genes were present in all DES isolates, while four were present in all but one genome (SAMEA7025205) (Supplementary Table 5). The final 45 putative resistance-associated genes included orthologs of *tetA(58)*, *abeS*, *aac(3)-VIIIa*, and *aph(6)-Ic*, potentially mediating resistance to tetracyclines, macrolides, and aminoglycosides. In addition to the intrinsic AxyXY-OprZ and AxyABM efflux systems, DES harbored eight additional RND efflux pump orthologs: six of *Pseudomonas aeruginosa* (including MexAB-OprM, MexCD-OprM, MexHI-OpmD, MexJK-OprZ, MexVW, MuxABC-OpmB); one AdeFGH system related to *Acinetobacter baumannii*, as well as ParRS two-component regulatory system ortholog; orthologs of one MacAB-OprZ ABC transporter and two MFS transporters (Figure 3).

**Figure 3.**
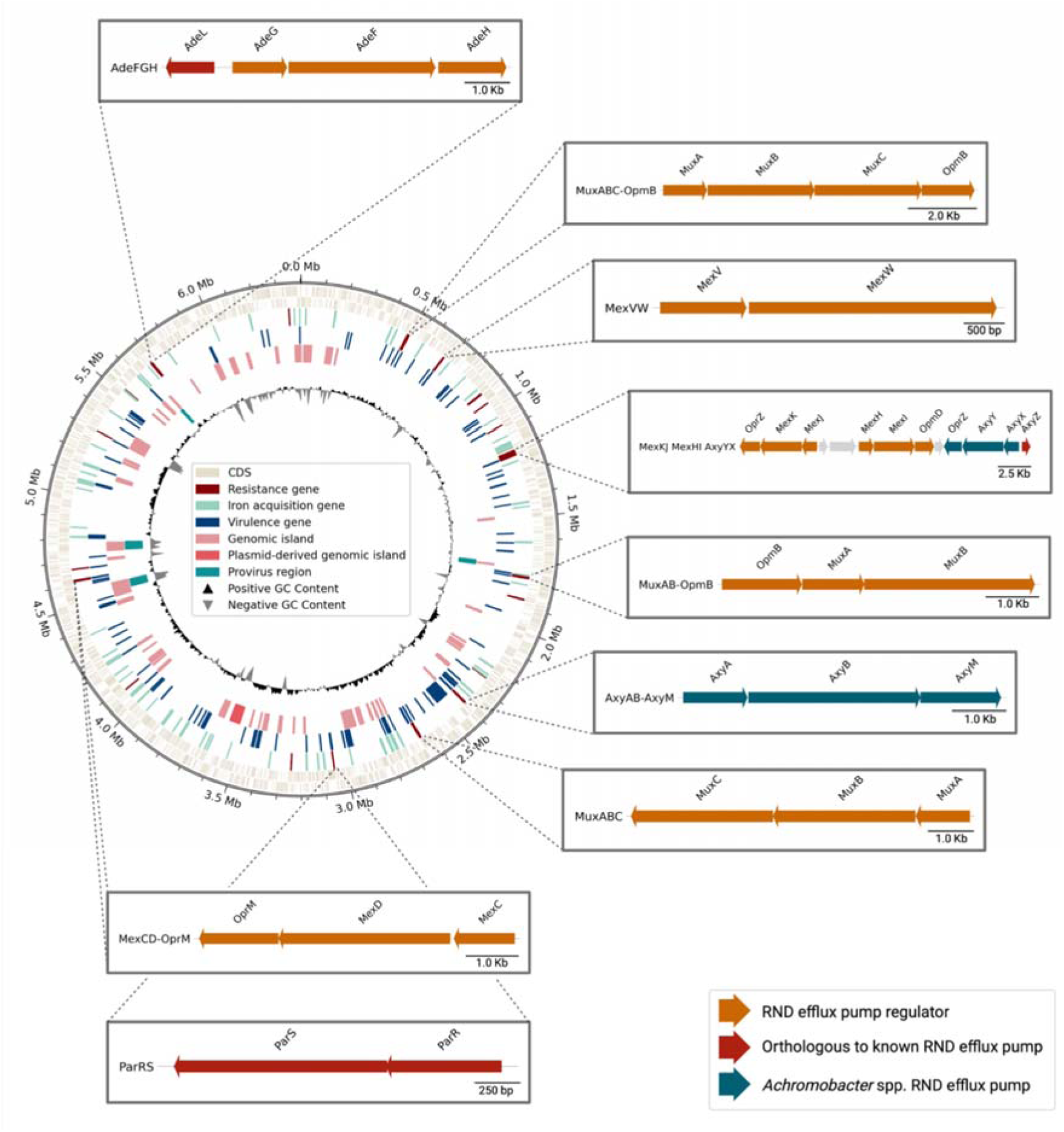
Circular representation of the DES reference genome (SAMEA7025168) highlighting genomic features associated with resistance, iron acquisition, and virulence. Predicted plasmid-like regions, prophage elements, and genomic islands are indicated together with the chromosomal location of RND efflux pumps.

When we examined the presence of these genes across all 139 *A. ruhlandii* genomes, most resistance determinants were conserved. Only two efflux systems showed variation: the MuxAB-OprM ortholog was absent in 16 genomes, while the MuxABC-OpmB ortholog was detected exclusively in DES isolates. In addition, two genomes lacked *axc-3*, *tet(A)* was absent in 31 genomes, and *tetA(58)* was present only in DES and absent from all non-DES isolates.

### Genomic islands, virulence, and iron-acquisition genes of DES

To characterise the DES genomic features we first identified genomic islands, known defence, virulence, and iron acquisition genes in the SAMEA7025168 genome that was used as a reference in downstream analyses (Figure 3).

Using PlasmidFinder, we detected an IncQ2_1 replicon within the region spanning 3,531,625–3,584,075, which was independently predicted by PIPdb as a plasmid-derived region. The region contained hallmarks of conjugative plasmids, including the *repC* gene and plasmid maintenance genes. However, the region was not located near repeated sequences or tRNA genes, which are common integration sites for integrative conjugative elements (ICEs)^81^. Therefore, we could not conclude that this region represents an ICE. Together, these findings support a plasmid origin of this chromosomal locus, suggesting that it is a plasmid-derived genomic island that is unique to DES and was not found in other isolates. We found that this sequence carries numerous genes possibly enhancing bacterial fitness: genes for plasmid transfer and maintenance (N=10); transcriptional regulators (N=14); stress and defense systems (N=10), including toxin–antitoxin modules, restriction-modification systems, putative heavy metal and aminoglycoside resistance genes; and one iron acquisition gene (Supplementary Table 6), suggesting that the plasmid-derived genomic island may have contributed to the adaptation, persistence, and antibiotic resistance of DES (Figure 3).

We predicted 65 genomic islands across the DES genome, encompassing 763 genes with a median of seven genes (range 1–51) (Supplementary Table 5). These genomic islands carried a substantial fraction of genes required for bacterial survival in the human environment, including 37 of the 128 iron-acquisition genes, 13 virulence genes, and 13 defense-related (COG V) genes. Across the full genome, we identified 178 virulence genes, dominated by genes involved in motility (N=68), nutritional and metabolic factors (N=41), effector delivery systems (N=25), and regulatory functions (N=13). Finally, we predicted four prophage-derived regions, each spanning 13–50 genes (Figure 3; Supplementary Table 5), adding another layer of mobile genetic content that may influence adaptation during chronic infection.

### DES likely emergence in 1980s

We estimated the time of emergence of the most common ancestor of DES using four independent approaches: BactDating, BEAST X, TimeTree and TempEst. All methods produced consistent estimates, placing the most likely emergence of DES between 1985 and 1989 (Figure 4). Root-to-tip regression analyses supported the presence of a temporal signal, with slopes of 3.78×10^-6^ substitutions/site/year (R²=0.40; s/s/y) in TreeTime and 3.91×10^-6^ s/s/y (R² = 0.41) in TempEst while BactDating estimated a mean evolutionary rate of 4.10×10^-6^ s/s/y (95% highest posterior density (HPD) 3.34×10^-6^–4.84×10^-6^). We obtained similar root age estimates across BEAST X models using different clock models and tree priors, further supporting the robustness of the emergence window with 95% HPD interval spanning from 1983 to 1993 and the average evolutionary rate of 4.45×10^-6^ s/s/y (95% HPD 4.02×10^-6^–4.89×10^-6^ s/s/y). These estimates are consistent with the first recorded DES isolate, collected in 1993, which was not available for inclusion in our phylogenetic analysis ^82^.

**Figure 4.**
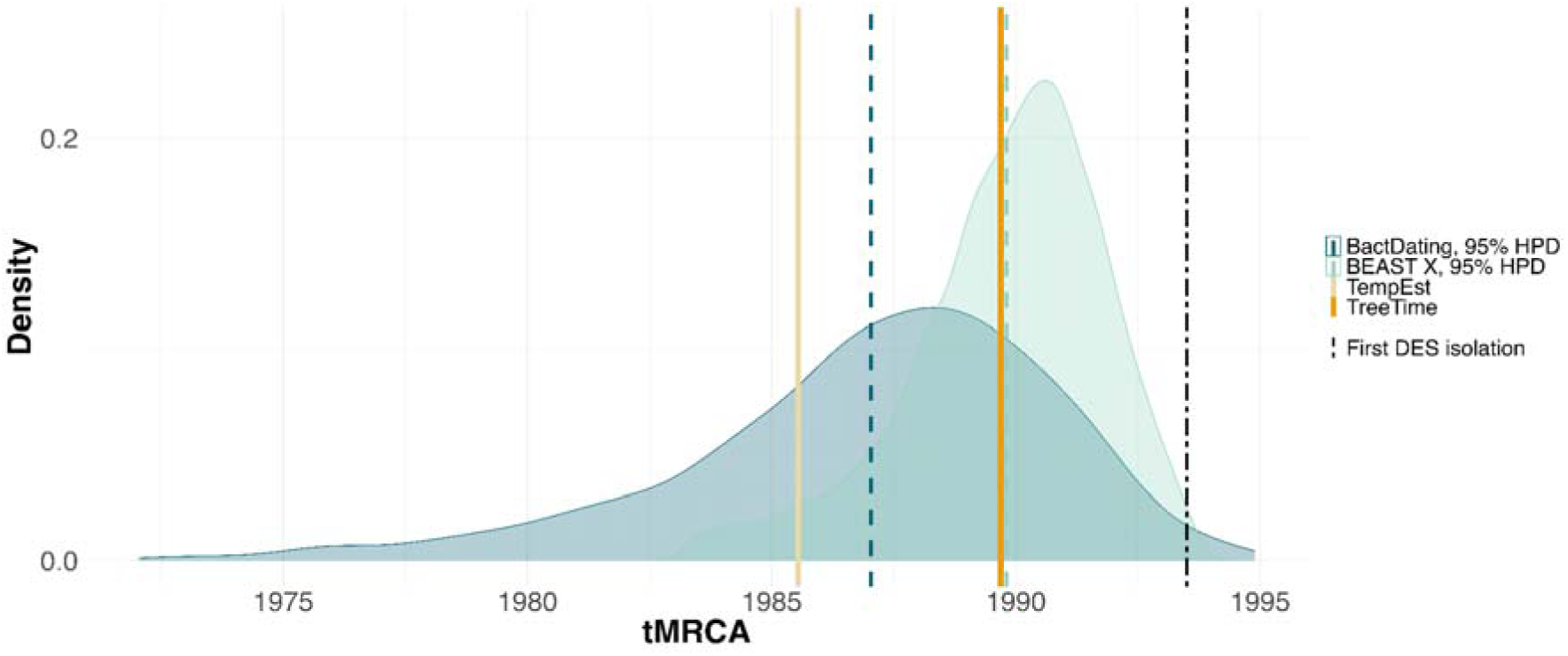
Estimates of the time to the most recent common ancestor (tMRCA) obtained using BactDating, BEAST X, TempEst, and TreeTime. BactDating and BEAST X display 95% highest posterior density (HPD) intervals, with the mean indicated by a coloured dotted line. The black vertical line marks the earliest known DES isolate, which was not included in the analysis.

### Recombination, regulation, and iron scavenging define the DES phenotype

Genome-wide association study (GWAS) was performed to identify genetic content differences between DES and non-DES isolates of *A. ruhlandii*. For this analysis we used a gene presence-absence matrix based on the DES reference genome. Consequently, the analysis was restricted to genes present in the DES reference and could not identify genes absent from the reference but present in non-DES isolates. A total of 1,302 tests were performed of which 279 genes were significantly associated with DES after Bonferroni correction and 194 of these genes were unique to DES (Supplementary Table 7; Figure 5).

**Figure 5.**
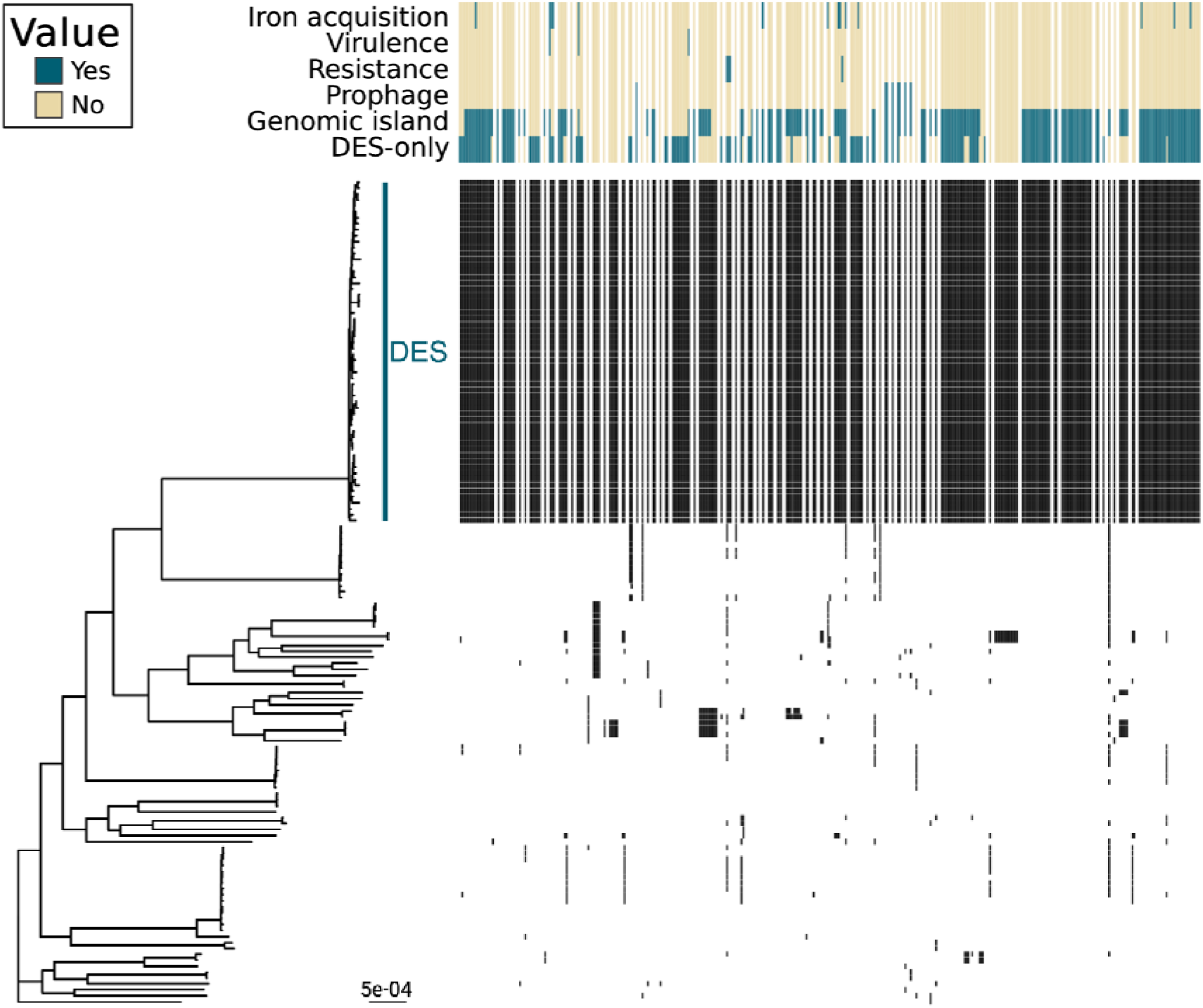
279 significant genes from genome-wide associations mapped onto the *A. ruhlandii* phylogeny. The left panel shows the phylogeny of *A. ruhlandii* isolates reconstructed from whole-genome alignment to the DES reference; branch lengths represent substitutions per site. The right panel displays a gene presence–absence matrix of loci identified in the GWAS analysis. Genes are ordered by their genomic position in the reference genome; adjacent associated genes are displayed without gaps, whereas gaps represent genomic intervals separating non-adjacent associated genes. Functional annotations and predicted gene origins are indicated above the matrix.

DES-associated genes clustered into discrete genomic regions, predominantly within genomic islands, suggesting extensive horizontal gene transfer (Figure 5). These regions were enriched for iron acquisition genes (11%; 14 of 128 total), highlighting iron scavenging as a potential adaptive trait. 12 of these iron acquisition genes were unique to DES (Supplementary Table 7). Genes involved in defense, replication, recombination and repair, and genes of unknown function were over twofold enriched among DES-associated loci, identifying recombination, resistance-related functions, and uncharacterised genes potentially associated with horizontally acquired genomic regions as major drivers of the DES phenotype. In contrast, genes involved in core cellular metabolism and housekeeping functions showed lower than expected enrichment, indicating that epidemic success is driven by accessory functions rather than changes to the core cellular functions (Figure 6).

**Figure 6.**
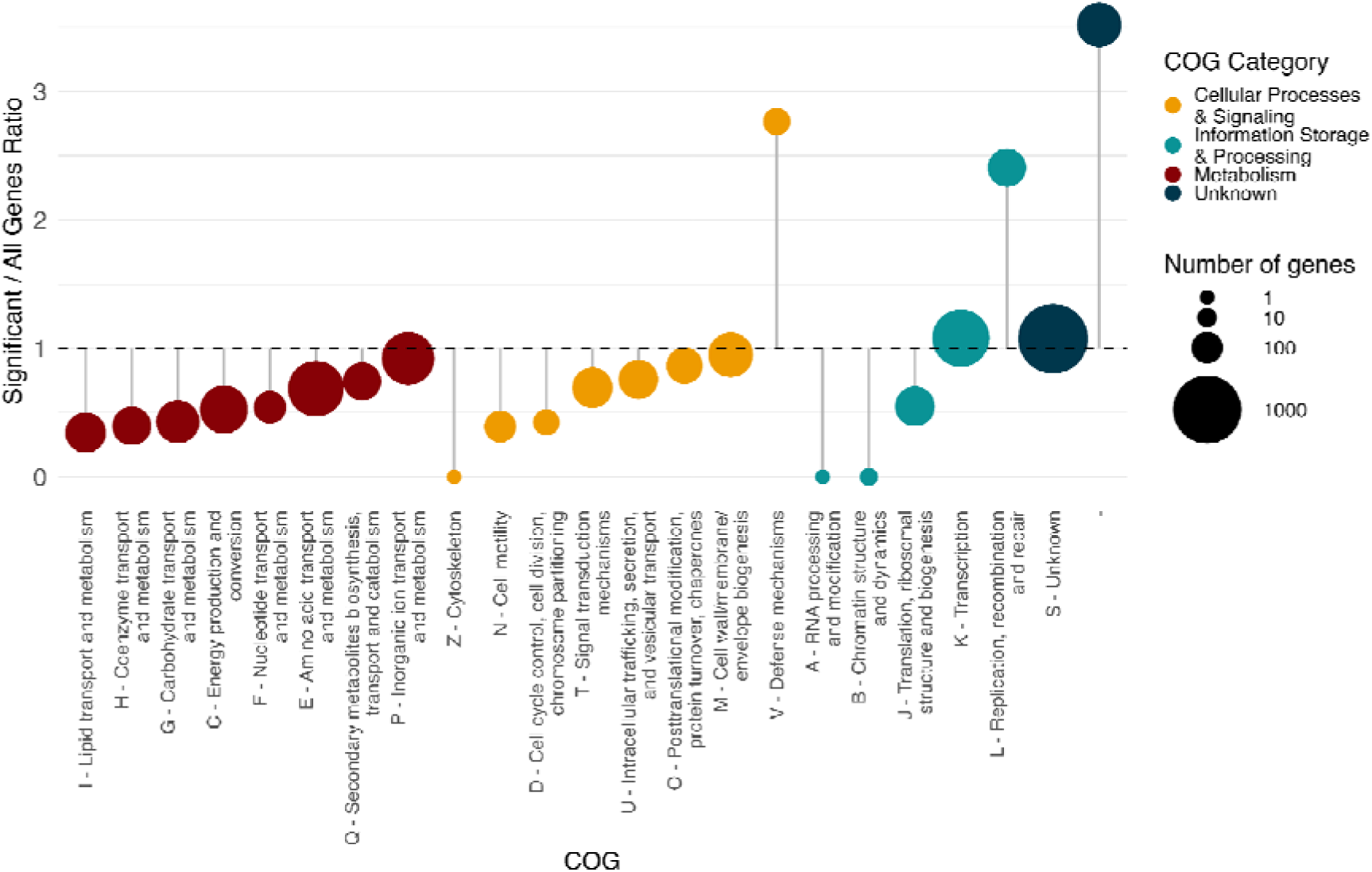
Ratio of gene functions that were significantly associated with DES during the GWAS. The horizontal dashed line represents the expected ratio between the associated genes and all of the genes in the DES reference genome.

Only three of the 178 annotated virulence genes showed significant association with DES, suggesting that known virulence factors do not contribute to the epidemic phenotype. In contrast, the *tetA*(58) gene and orthologs of the MuxABC RND efflux pump were only observed in DES genomes, suggesting DES-unique enhanced multidrug resistance as a contributor to the epidemic phenotype (Figure 5; Supplementary Table 7).

A MetaGraph ^79^ search against 2.7 million bacterial assemblies from AllTheBacteria ^23^ database found that 71% (139/194) of the predicted genomic island regions were not detected in any bacterial genomes outside DES. In contrast, 77% (65/85) of the significantly associated genes located outside genomic islands had sequence matches in other *Achromobacter* spp. isolates (Supplementary Table 7). Furthermore, 69% (38/55) of genes with no COG function assigned (COG -) were not identified in any other bacterial assemblies (Supplementary Table 7) .

### Continuous DES evolution

Using BEAST X under a constant population size model, we estimated the molecular clock rate and found that the evolutionary rate of DES is consistently high across the lineage (Supplementary Figure 4). Using the coalescent Skygrid tree prior, we examined effective population size changes since the emergence of DES. The Skygrid profile shows a consistently positive effective population size early in the lineage, indicating population expansion, followed by apparent stabilisation in recent years (Supplementary Figure 5). DES exhibited the highest dN/dS ratio of 1.19, compared with 0.91 for US-ES and 0.68 for ST36. The elevated dN/dS ratio in DES reflects a greater relative contribution of non-synonymous substitutions, while in US-ES and ST36 it is consistent with purifying selection.

### Comparable genomic features across independent epidemic *A. ruhlandii* lineages

Beyond DES, we identified two additional epidemic *A. ruhlandii* lineages, using the definition of an epidemic strain as a genetically distinct lineage that undergoes prolonged transmission across individuals or populations in more than one geographic location. The first corresponds to sequence type ST36, which originated in Russia and has subsequently been detected in the United Kingdom and the United States during more than a decade of transmission. The second represents a distinct United States epidemic strain (US-ES), which was isolated across three different US states over a nine-year period (Supplementary Table 2; Figure 7).

**Figure 7.**
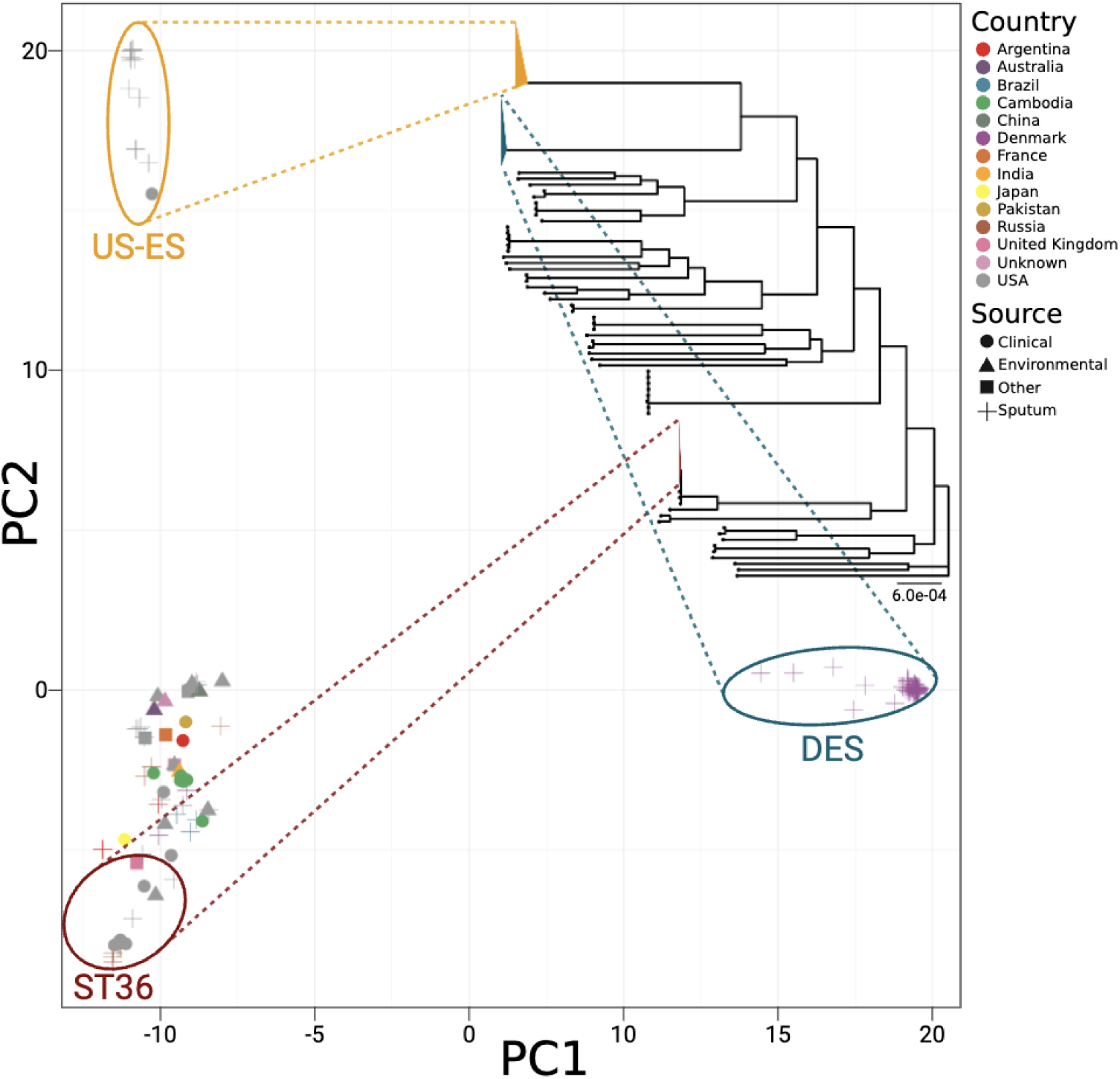
Principal component analysis of the core *A. ruhlandii* genome multiple sequence alignment with epidemic strains highlighted and connected to the core-genome-based phylogenetic tree. The phylogeny was rooted using *A. dolens* as an outgroup, which was subsequently removed from the displayed tree. Branch lengths represent substitutions per site.

To identify if similar genomic features are associated with the epidemic phenotype, we associated gene presence-absence from pangenomic analysis of the three epidemic lineages with non-epidemic *A. ruhlandii* genomes. In total, we associated 61 (DES), 105 (ST36) and 204 (US-ES) genes with epidemic lineages. 0–4 genes were associated with epidemic phenotypes between the three datasets (Figure 8). Similarly to DES, we observed genes of unknown function and lacking annotation (COG=S/–; N=13 in DES; N=52 in ST36; N=98 in US-ES; Supplementary Table 8) and transcription genes (COG=K; N=10 in DES; N=12 in ST36; N=27 in US-ES; Supplementary Table 8) being associated with epidemic phenotypes (Figure 8). Enrichment for iron acquisition genes was specific only to DES with 10% of all associated genes annotated as related to iron acquisition; however, it was not observed in other ES at ∼3% of associated genes (N=6 in DES; N=3 in ST36; N=6 in US-ES; Supplementary Table 8). Furthermore, DES was confirmed to carry a 12 amino acid deletion in *mutS* gene (Supplementary Table 2) and exhibited a markedly elevated transition-to-transversion (Ts/Tv) ratio, consistent with its hypermutator phenotype, with an estimated ratio of 24.34 (95% HDR, 22.80–26.02). This was substantially higher than those observed in the other ESs (US-ES [6.12; 95% HDR, 5.12–7.13] and ST36 [2.72; 95% HDR, 2.35–3.11]), as well as in non-ES isolates (5.09; 95% HDR, 5.06–5.13).

**Figure 8.**
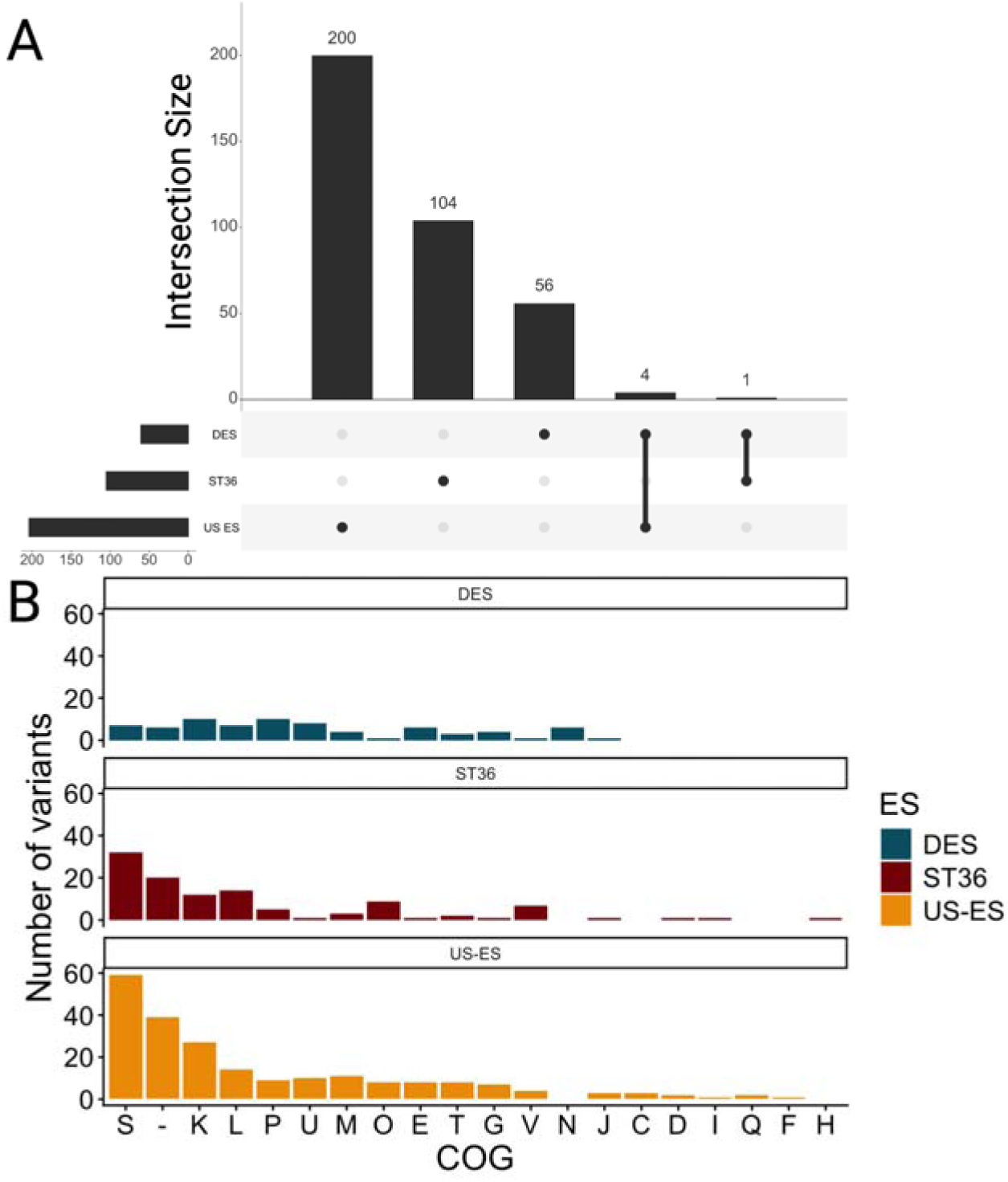
Epidemic phenotype and gene associations across the three epidemic *A. ruhlandii* lineages (DES, US-ES, and ST36). (A) UpSet plot showing shared and unique genes across the epidemic lineages. (B) The distribution of associated genes broken down by COG functional categories.

### Epidemic strain pangenome exploration

We constructed the *A. ruhlandii* pangenome which comprised 11,929 genes: 3,619 hard-core, additional 1,152 soft-core, and 7,158 accessory genes. This pangenome framework was then used to compare the distribution of iron acquisition, virulence, and resistance genes among DES, other epidemic strains (ES), and non-epidemic isolates.

Across the pangenome, we identified 85 virulence-associated, 111 resistance-associated, and 166 iron acquisition–associated genes (Supplementary Table 9). DES was enriched in iron acquisition genes relative to non-DES lineages, with an estimated increase of 7.18 genes (95% HDR 0.08–14.22; Supplementary Figure 6). No comparable enrichment was observed for virulence or resistance genes (95% HDR -7.69–2.01 and -2.74–6.52, respectively; Supplementary Figure 6). Notably, relative to non-DES lineages, DES carried an estimate of 1.99 fewer effector delivery system–associated virulence genes but 2.68 more abundance of RND efflux system genes (95% HDR -7.40–0.51 and -0.11–5.39, respectively; Figure 9, Supplementary Figure 6). This increase in RND efflux genes was driven by a DES-unique MuxABC RND efflux pump ortholog. These patterns remained consistent when DES was compared against non-ES strains rather than non-DES strains (Supplementary Figure 7). When all three epidemic strains (DES, US-ES, and ST36) were analysed jointly, epidemic lineages carried fewer virulence genes than non-ES strains, with an estimated reduction of 1.34 virulence genes overall, including 2.24 fewer effector delivery system genes (95% HDR -5.75 – -0.44 and -4.18 – -0.33, respectively; Supplementary Figure 7). In contrast, iron acquisition and resistance gene content did not differ substantially between epidemic and non-epidemic strains, underscoring the distinct enrichment of iron acquisition genes observed specifically in DES (Supplementary Figure 7; Figure 9).

**Figure 9.**
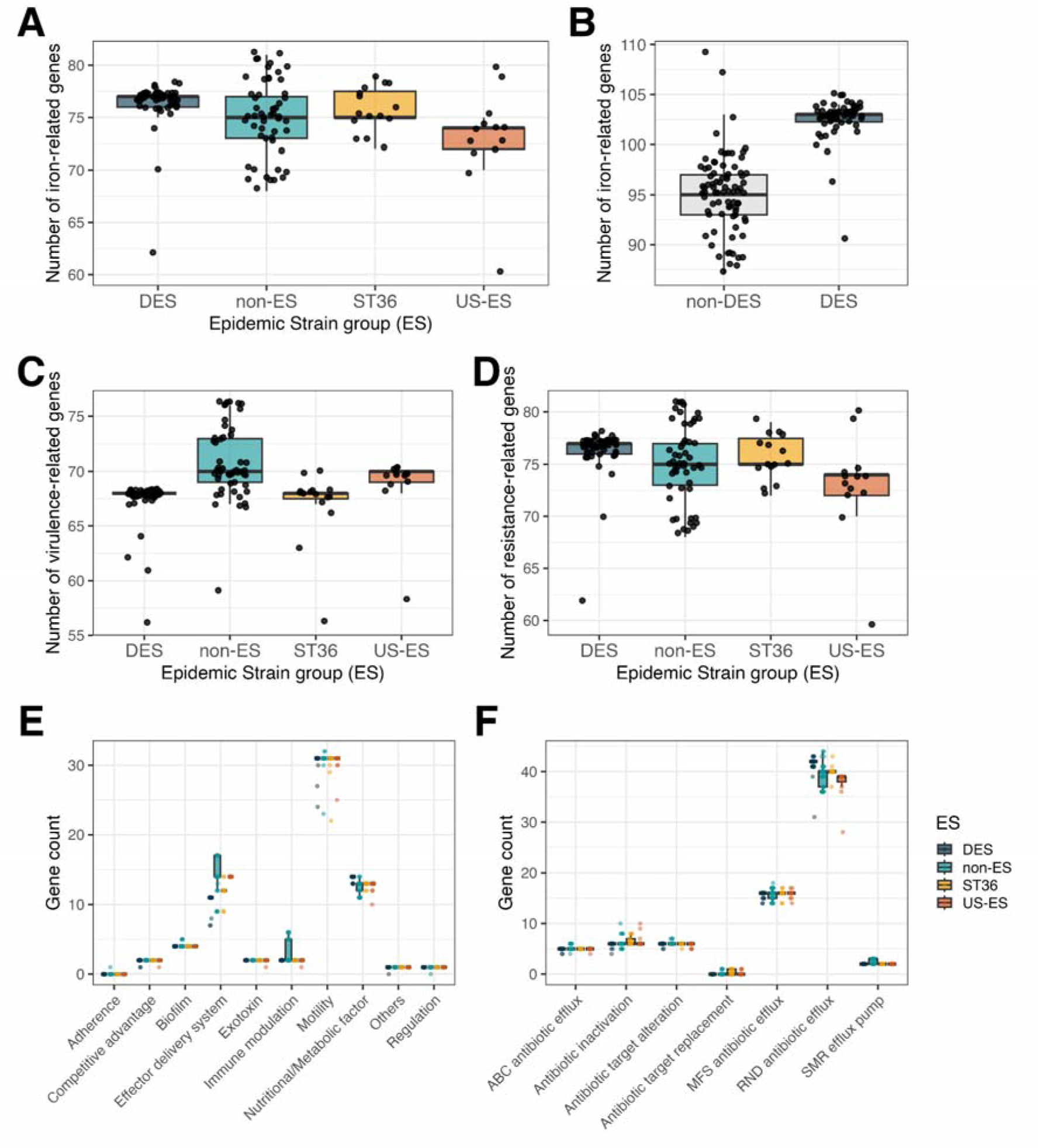
Enrichment for iron acquisition, resistance, and virulence gene content across DES and other *A. ruhlandii* isolates. Panels show enrichment of (A-B) iron acquisition genes, (C) virulence genes, further subdivided by (D) effector delivery system–associated virulence factors, and (E) resistance genes, further stratified by (F) resistance mechanism.

## Discussion

*Achromobacter* spp., particularly epidemic lineages such as DES, are increasingly recognised as clinically relevant opportunistic pathogens in pwCF ^2,83^. At the same time, a growing body of evidence shows that opportunistic environmental bacteria, including *Achromobacter* spp., are emerging as causes of infections in other vulnerable populations, such as individuals injured in war zones or neonates in intensive care settings ^5,84^. Understanding how these bacteria evolve to adapt to host environments and give rise to epidemic lineages is therefore becoming increasingly important. The DES lineage provides a rare opportunity to investigate the evolutionary processes underlying the emergence and persistence of an epidemic bacterial strain in real time.

DES appears to be confined to Denmark, with no confirmed occurrence elsewhere. This pattern is consistent with the CF-associated epidemic *P. aeruginosa* lineages DK01 and DK02 that, likewise, have not been detected outside Denmark ^85,86^. By contrast, the additional epidemic *A. ruhlandii* lineages ST36 and US-ES identified here have persisted for over a decade and have been isolated across multiple countries or U.S. states. The broader global distribution and spread of epidemic *A*. *ruhlandii* lineages remain difficult to quantify as the genus is still markedly undersampled ^5,87^.

Our phylogenetic analyses indicate that DES’ most recent common ancestor likely emerged in the late 1980s. This estimate is supported by the monophyletic DES clade, the earliest recorded isolate from 1993 ^82^, and consistent results across multiple molecular clock methods, suggesting it reflects the true origin of the DES lineage. Previous studies have attributed the DES hypermutator phenotype to a 12-amino-acid deletion at positions 174–185 of the *mutS* gene ^9,82^. Our genomic analysis confirmed the presence of this deletion across the DES lineage, supporting the previously described genetic basis of its hypermutator phenotype. This phenotype was further reflected in the substantially higher Ts/Tv ratio observed in DES than in other ESs and non-ES isolates further supporting the hypermutator phenotype of DES lineage. We also show that DES has maintained a consistently high evolutionary rate throughout the lineage existence, indicating ongoing rapid accumulation of genetic variation that may facilitate continued adaptation to the host environment ^9,88^.

Together, our findings support a model in which DES originated from an environmental *A. ruhlandii* lineage and subsequently underwent continuous rapid adaptation to the CF airway environment, characterised by extensive genomic diversification, accumulation of nonsynonymous mutations, and acquisition of host-associated traits, ultimately facilitating patient-to-patient transmission and epidemic spread. However, the absence of closely related non-DES or environmental isolates prevents direct testing of this evolutionary scenario.

The inferred effective population size increased following the emergence of DES is consistent with an initial lineage expansion, confirming DES’ epidemic nature. Notably, no DES transmission events were detected in the last two years, suggesting that the effective population size is likely to decline further and that circulation of this lineage among pwCF could potentially be eliminated. This trend is plausibly associated with the introduction of triple-combination CFTR modulator therapy, which reduces bacterial burden and antibiotic exposure in many patients, thereby limiting opportunities for transmission and within-host adaptation ^18^. However, accumulating evidence indicates that CFTR modulator therapy does not result in complete eradication of chronic infections ^89,90^, implying that the risk posed by highly host-adapted lineages such as DES is reduced but not eliminated.

DES shows a gradual decrease in antibiotic resistance over time. While this trend is an anticipated consequence of CFTR modulator therapy, similar effects have not been consistently observed across studies of CF airway pathogens ^91–93^. This apparent decline may be influenced by confounding factors, particularly the marked reduction in DES transmission in recent years. Nonetheless, our dataset spans up to four years post-introduction of CFTR modulators and includes a larger sample, potentially capturing effects that earlier studies were unable to detect ^91,93,94^.

Despite this decrease, DES retains a broad resistome, and currently known resistance determinants, including multidrug efflux systems, β-lactamases, and genes acquired through horizontal gene transfer, do not fully account for the high resistance levels observed in this lineage ^1,13,14^. Among these are two previously characterised RND efflux systems, AxyXY-OprN and AxyAB-OprM that confer resistance to aminoglycosides, β-lactams, fluoroquinolones, and chloramphenicol ^95,96^. In addition, we identified nine putative RND efflux systems and multiple other resistance-associated genes, indicating that *A. ruhlandii* genomes harbour a number of RND efflux pumps comparable to that of *P. aeruginosa*, where such systems are key for the development of multidrug resistance ^97^. As efflux-mediated resistance depends strongly on gene expression and regulatory context, genomic presence alone does not reliably predict resistance phenotypes ^98,99^. Nevertheless, the high density of RND efflux systems across *A. ruhlandii* genomes—of which only one putative system is unique to DES, with the remainder present in the majority of isolates—points to a substantial intrinsic capacity for resistance evolution and underscores the potential for the emergence of new epidemic lineages. Similar abundance of RND efflux pumps has been reported in other *Achromobacter* species, suggesting that this feature is not unique to *A. ruhlandii* ^13,83^.

Notably, several recently described RND efflux pumps in other *Achromobacter* spp. cannot yet be assessed here because their sequences are not publicly available ^100^.

To identify genomic features distinguishing epidemic from non-epidemic lineages, we performed GWAS analyses of the gene content, which showed that DES differed from non-DES isolates primarily through enrichment of genes of unknown function (COG S) and genes without functional annotation (COG -), many of which were not detected in other bacterial genomes and were associated with mobile genetic elements. This pattern is consistent with extensive horizontal gene transfer during lineage emergence and is further supported by the significant enrichment of recombination-related genes in DES, as well as by observations from genomic studies of epidemic strain emergence in other bacterial species. ^101,102^. Notably, our GWAS was based on gene presence–absence, it does not capture associations arising from sequence variation within shared genes. Collectively, our findings indicate that horizontal gene transfer has been a major driver of DES evolution. The integration of plasmid-origin genomic island may have contributed to the epidemic phenotype, although its precise role remains unclear; nonetheless, it aligns with the broader trend of acquiring novel genomic content during lineage emergence.

In contrast, GWAS revealed negative enrichment of core cellular metabolism and housekeeping genes, indicating that variation in these essential functions is likely selected against, presumably because it compromises basic cellular fitness. This pattern suggests that epidemic success is driven less by rewiring core metabolism and more by expanding accessory functions that enhance adaptability and persistence. Notably, the order in which these gene clusters were acquired and their individual contributions to the epidemic phenotype cannot be determined, as no intermediate isolates representing the evolutionary transition between non-DES *A. ruhlandii* and DES are available.

One of the most striking features of DES is its markedly higher iron acquisition gene content compared with non-DES lineages. In humans, iron is tightly sequestered by host proteins and is therefore not freely available to invading bacteria. Efficient iron scavenging in this iron-limited environment is a key determinant of bacterial growth, persistence and successful host colonisation, particularly during chronic infection ^103^. The enrichment of iron acquisition systems in DES therefore highlights iron scavenging as a likely contributor to its epidemic success, consistent with known mechanisms of host adaptation in *Pseudomonas aeruginosa* and *Vibrio* spp. ^103,104^. This interpretation is further supported by recent evidence showing increased siderophore-mediated iron acquisition in clinical CF *Achromobacter* isolates, with levels comparable to those observed in CF-adapted *P. aeruginosa* strains ^105^. In addition to enhanced iron acquisition capacity, DES also carries a higher number of putative RND efflux pump genes, in particular, an ortholog of the MuxABC RND efflux system, which in *P. aeruginosa* confers resistance to aztreonam, novobiocin, tetracycline, erythromycin, kitasamycin, and rokitamycin ^106^. This expanded efflux capacity may contribute to the successful establishment and persistence of DES, although other epidemic strains do not show the same enrichment of resistance genes, indicating that this is not a universal feature of epidemic strains.

Beyond gene content, DES exhibits hypermutability and an increased accumulation of nonsynonymous mutations. Hypermutability is a known feature of chronic, host-adapted infections and is expected to accelerate within-host adaptation ^14,107^. However, comparable mutational patterns were not observed in the other epidemic lineages (ST36 and US-ES), indicating that hypermutation is not a prerequisite for epidemic success but may instead enhance persistence and adaptive potential once a lineage is established. Including additional epidemic lineages in the analysis revealed a set of features shared across epidemic strains. Notably, epidemic lineages exhibited a reduced number of virulence factor genes, particularly those associated with effector delivery systems. This reduction may reflect selection for lower immunogenicity, facilitating long-term persistence within the host by decreasing immune activation ^108,109^.

Taken together, these findings indicate that DES represents a highly dynamic epidemic lineage. Its emergence and persistence appear to reflect a combination of extensive horizontal gene transfer, substantial intrinsic resistance potential, increased iron acquisition capacity, reduced effector-mediated virulence, and an accelerated accumulation of nonsynonymous variation. Analyses of additional epidemic lineages allow some features of epidemic *Achromobacter* strains, such as enrichment of genes of unknown function and reduction of virulence genes, to be generalised, while others, including iron acquisition enrichment and hypermutability, appear to be DES-specific. Collectively, these observations suggest that *A. ruhlandii* possess an intrinsic capacity to repeatedly evolve epidemic lineages, with a pre-existing high resistance potential providing a genomic background that can facilitate this transition under appropriate selective pressures, such as high antibiotic exposure or infection in highly vulnerable environments, including war-associated injuries and hospital acquired infections ^5,84^.

## Conclusions

Our results indicate that *A. ruhlandii* has a high intrinsic potential for repeated emergence of novel epidemic strains beyond those currently recognised. In DES, a combination of high intrinsic antibiotic resistance potential, reduced effector-mediated virulence, enhanced iron acquisition, and episodic recombination through plasmid and genomic island acquisition enabled rapid emergence, sustained patient-to-patient transmission, and long-term persistence.

These findings underscore the need for proactive surveillance strategies aimed at early detection of emerging epidemic lineages. Increased sampling density and routine integration of whole-genome sequencing with phenotypic susceptibility testing would allow timely identification of transmission, improved understanding of epidemic strain development, and more informed infection-control and patient management decisions. Such combined analyses are also essential for assessing how changes in treatment strategies, including CFTR modulator therapy, reshape the evolutionary dynamics of chronic airway pathogens.

## Acknowledgements

This work is supported by HORIZON-MSCA-2023-PF-01 from the European Commission (Grant Agreement No. 101151221). The authors are thankful for the HPC resources provided by the IT Research Center of Vilnius University; for the ONT sequencing support provided by UAB Seqvision. GD received support from the Research Council of Lithuania, Lithuania under the EMBO Installation Grant programme under grant No 5305.

## Author contribution

MG, HKJ, RLM, and GD conceptualised the study. MG performed the analyses and drafted the manuscript. All authors contributed to manuscript revision and approved the final version.

## Data availability

Raw sequencing data of 36 newly sequenced *A. ruhlandii* isolates are available on European Nucleotide Archive under PRJEB110520 BioProject study accession number.

## Supplementary data

**Supplementary Figure 1.** All clinical samples taken from people with CF that were infected with *Achromobacter* spp. and had isolates sequenced at least once between 2002 and 2024.

**Supplementary Figure 2.** Bioinformatics workflow flowchart representing all the analyses performed in this study.

**Supplementary Figure 3.** Susceptibility testing of all patients infected with *A. ruhlandii* in 2002–2024 with the two-year sliding window fraction of resistance to antibiotics (top) and the total count of *A. ruhlandii* samples during that period (bottom). Trendlines were fitted using locally estimated scatterplot smoothing (LOESS) for each patient (colored lines) and across all patients (gray line); the shaded area around the gray trendline represents the 95% confidence interval of the fitted overall LOESS trend.

**Supplementary Figure 4.** Estimated molecular clock rate across the Danish Epidemic Strain phylogeny. Branch colors and numbers on branches indicate the inferred molecular clock rate, with blue representing lower rates and red representing higher rates. The phylogenetic scale represents estimated node ages.

**Supplementary Figure 5.** Skygrid analysis of the Danish Epidemic Strain using the coalescent Skygrid tree prior showing the summary statistics and estimated value distributions (top) and a Skygrid reconstruction (bottom).

**Supplementary Figure 6.** Summary statistics of the association analysis between the Danish Epidemic Strain and iron (A), virulence (B), effector delivery systems (C), resistance (D), RND efflux pumps (E) using Bayesian generalised linear multivariate models implemented in the brms R package accounting for shared ancestry and repeated sampling from the same patient.

**Supplementary Figure 7.** Summary statistics of the association analysis between the Epidemic Strains and iron (A), virulence (B), effector delivery systems (C), resistance (D), RND efflux pumps (E) using Bayesian generalised linear multivariate models implemented in the brms R package accounting for shared ancestry and repeated sampling from the same patient.

**Supplementary Table 1.** Raw susceptibility information. R – resistant; R(Inh) – resistant, could be used for inhalations; I – susceptible, increased exposure; S – susceptible; NA – not tested/information not available.

**Supplementary Table 2.** A list of all 159 *A. ruhlandii* genomes together with the metadata, assembly method and assembly statistics.

**Supplementary Table 3.** Antibiotic-specific resistance trends among DES isolates from 2002–2024 estimated using generalised additive mixed models.

**Supplementary Table 4.** A list of genes identified in the reference DES genome using AMRFinderPlus together with their statistics.

**Supplementary Table 5.** Annotations of 5,880 reference DES genome genes with metadata defining the gene as being virulence, resistance, plasmid-derived genomic island, defense, iron acquisition and/or genomic island gene.

**Supplementary Table 6.** Gene annotations from the plasmid-derived genomic island sequence in a DES reference genome.

**Supplementary Table 7.** 279 with DES phenotype significantly associated genes and their annotations.

**Supplementary Table 8.** 370 with ES (US-ES, DES or ST36) phenotype significantly associated genes and their annotations.

**Supplementary Table 9.** Annotations of 10,225 *A. ruhlandii* pangenome genes with metadata defining the gene as resistance, virulence or iron acquisition gene.

