## Supplementary figures and images for "Thirty years of *Achromobacter ruhlandii* evolution reveal pathways to epidemic lineages"

### Supplementary figure 1

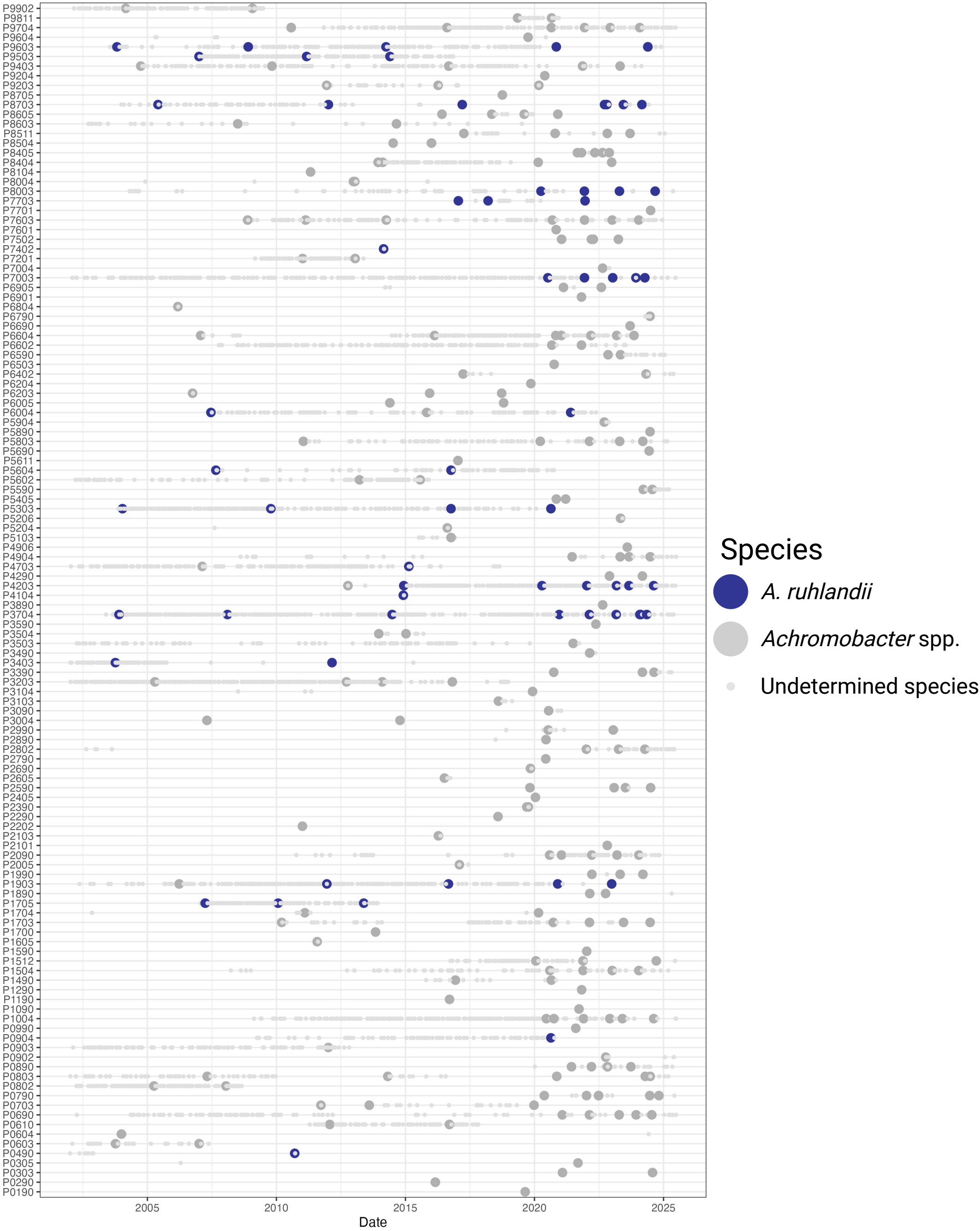

### Supplementary figure 2

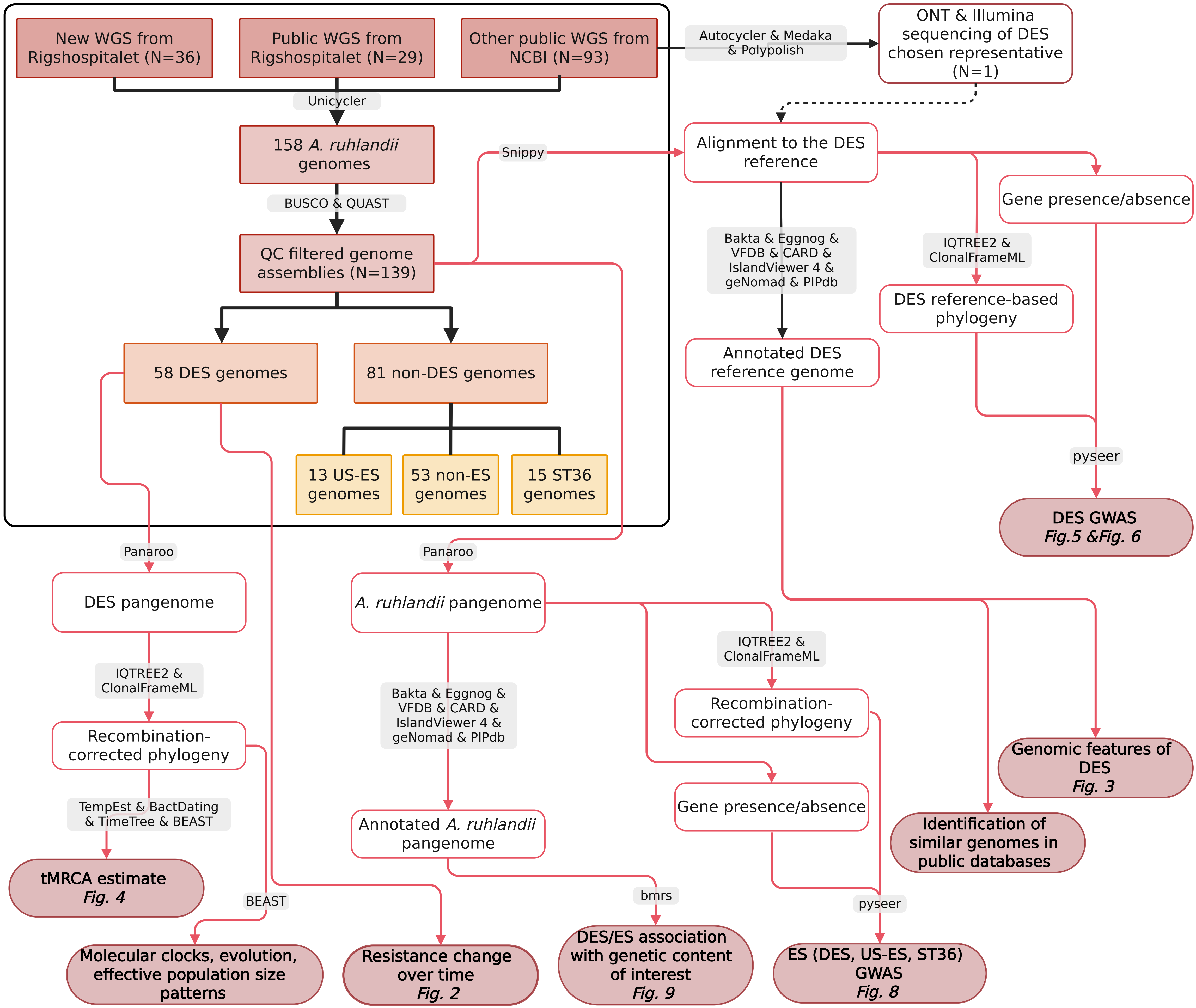

### Supplementary figure 3

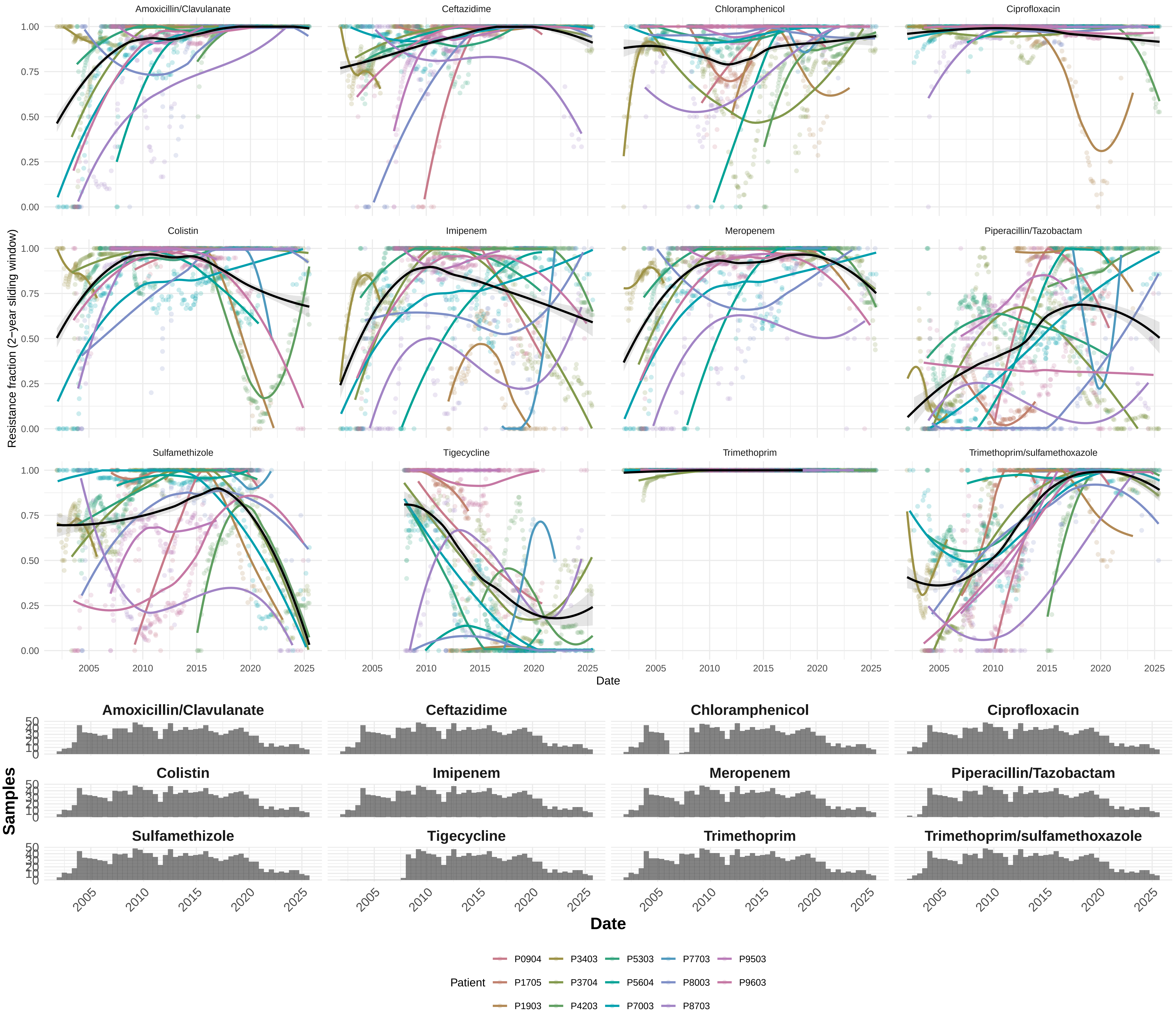

### Supplementary figure 4

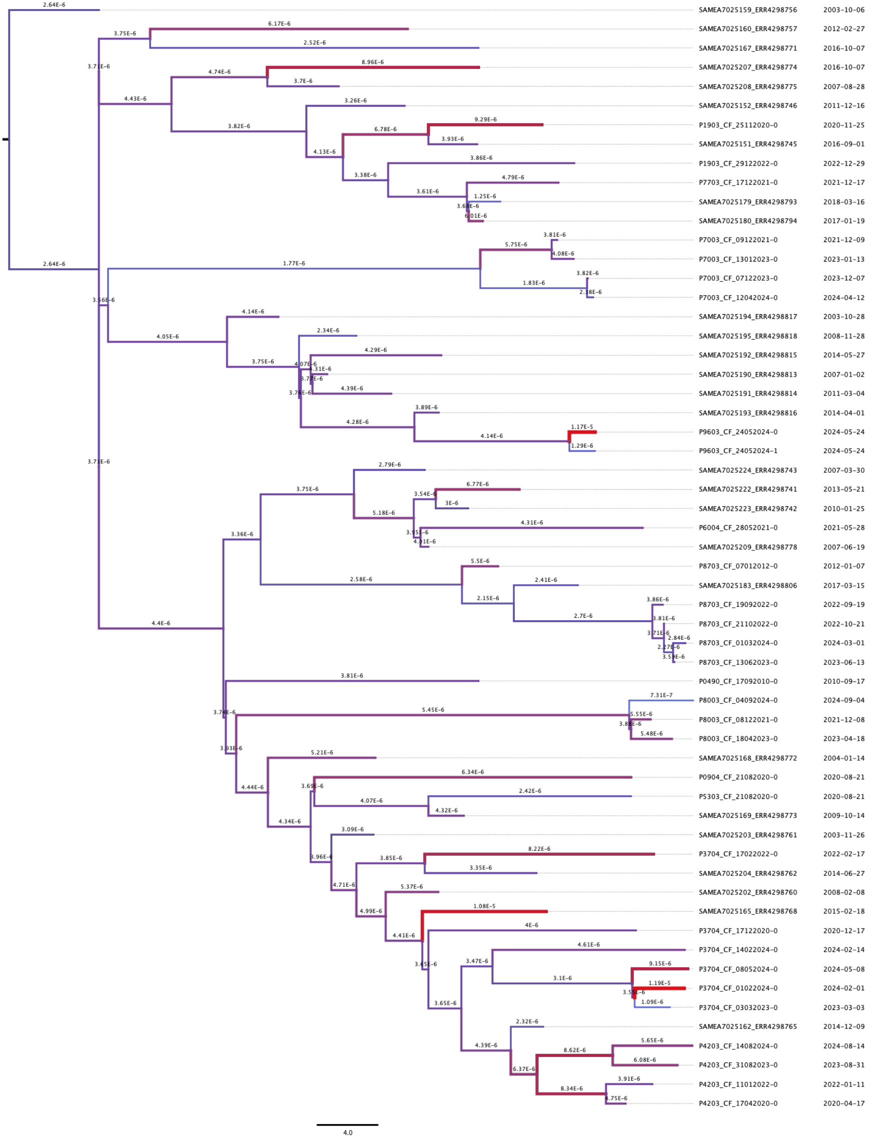

### Supplementary figure 6

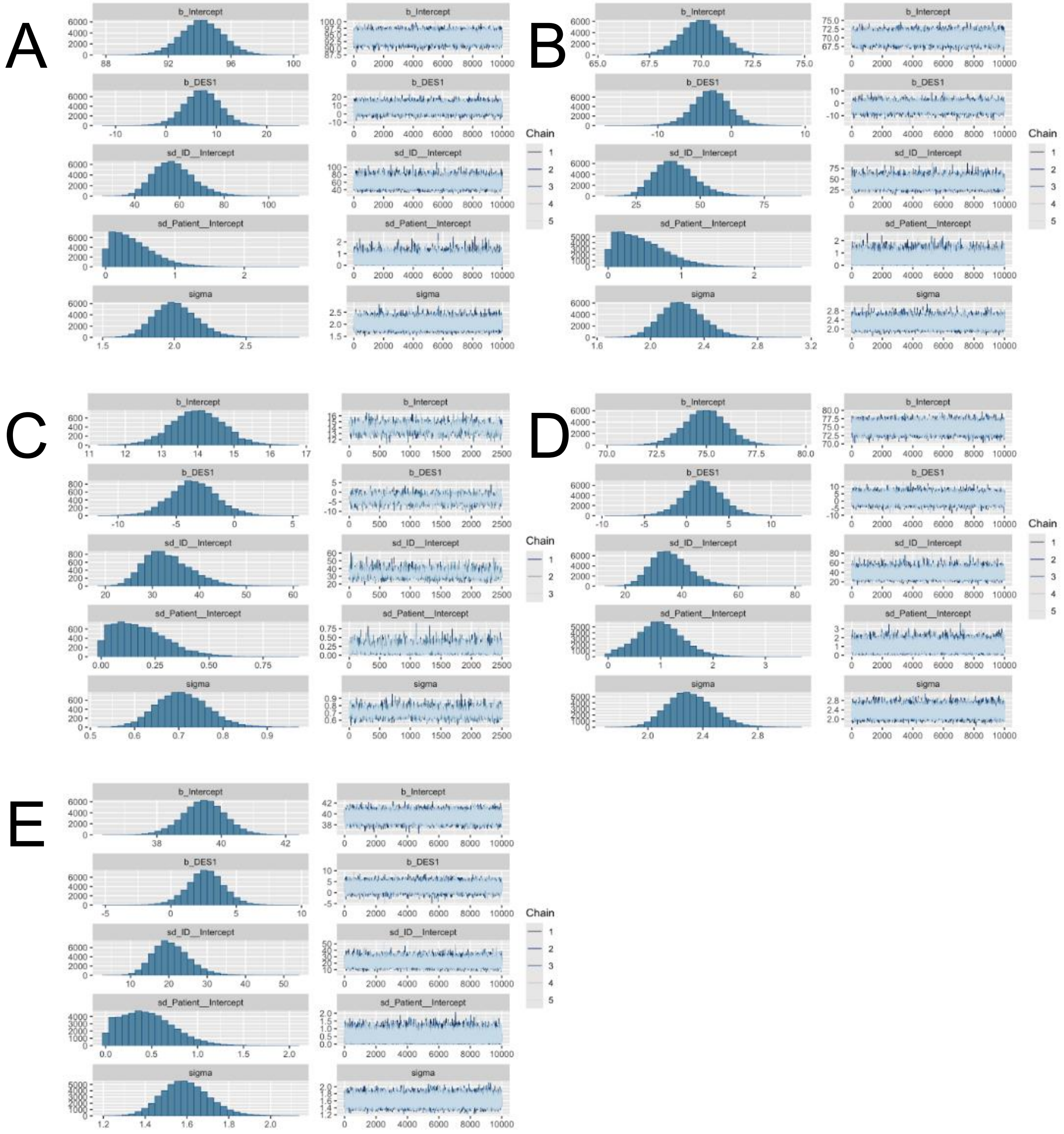

### Supplementary figure 7

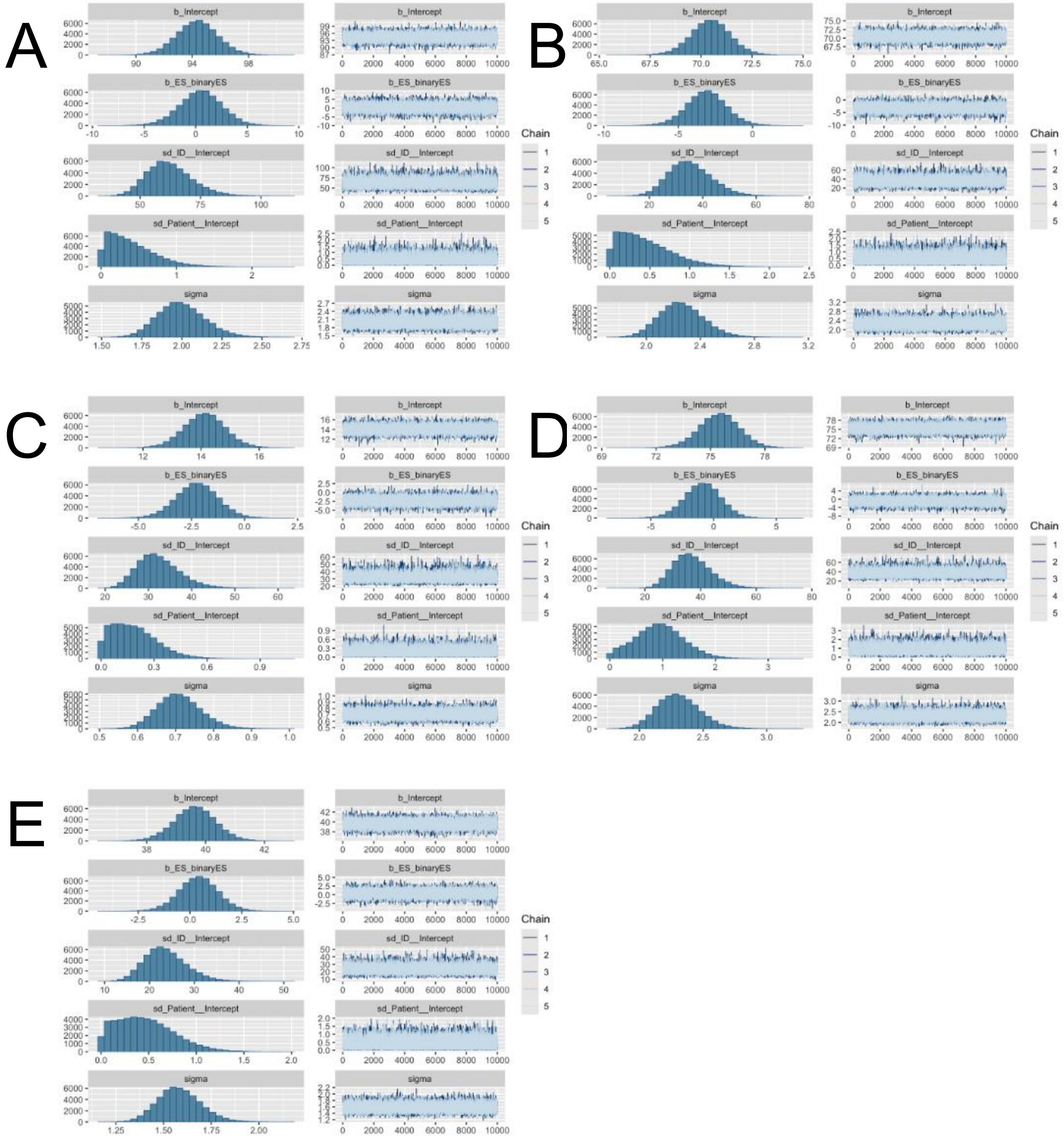
