## Supplementary figure 5 for "Thirty years of *Achromobacter ruhlandii* evolution reveal pathways to epidemic lineages"

Estimates | Marginal Density | Joint-Marginal | Trace

| Summary Statistic | skygrid.logPop... | skygrid.logPop... | skygrid.logPop... | skygrid.logPop... | skygrid.logPop... | skygrid.logPop |
|---|---|---|---|---|---|---|
| mean | 4.2351 | 5.2001 | 4.5104 | 4.0505 | 0.97 | 0.9345 |
| stderr of mean | 2.9606E-3 | 4.035E-3 | 2.7095E-3 | 7.1893E-3 | 0.0543 | 0.0375 |
| stdev | 0.2914 | 0.3482 | 0.2486 | 0.3453 | 0.7893 | 2.643 |
| variance | 0.0849 | 0.1212 | 0.0618 | 0.1193 | 0.623 | 6.9856 |
| median | 4.2221 | 5.1802 | 4.4987 | 4.0292 | 0.9428 | 1.0786 |
| value range | [3.3084, 5.5... | [4.1382, 6.9... | [3.6771, 5.6... | [2.9669, 5.7... | [-1.7762, 3.... | [-35.6785, 1 |
| geometric mean | 4.2251 | 5.1885 | 4.5036 | 4.036 | n/a | n/a |
| 95% HPD interval | [3.6857, 4.8... | [4.5447, 5.9... | [4.0349, 5.0... | [3.3938, 4.7... | [-0.5127, 2.... | [-4.3276, 5.. |
| auto-correlation time (ACT) | 6969.4608 | 9067.4104 | 8021.9764 | 29262.9457 | 3.1898E5 | 13613.4214 |
| effective sample size (ESS) | 9687.3 | 7445.9 | 8416.3 | 2307.2 | 211.7 | 4959.4 |
| number of samples | 13503 | 13503 | 13503 | 13503 | 13503 | 13503 |

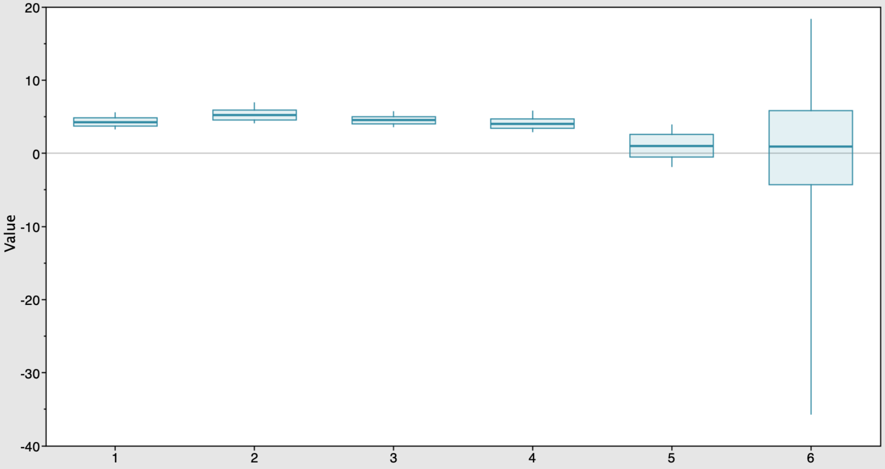


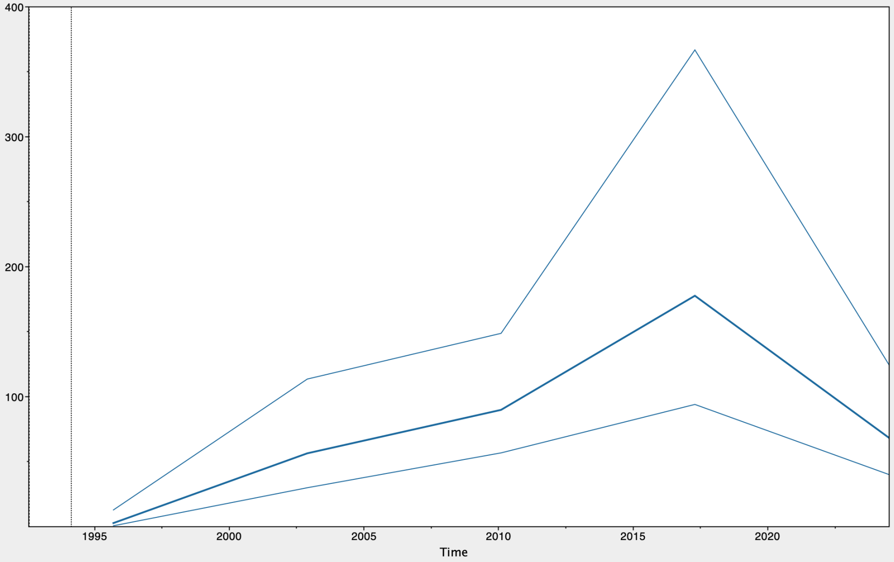
